# Perturbation-driven transcriptional heterogeneity impacts cell fitness

**DOI:** 10.1101/2024.05.31.596868

**Authors:** Mariona Nadal-Ribelles, Carme Solé, Anna Diez-Villanueva, Camille Stephan-Otto Attolini, Yaima Matas, Lars Steinmetz, Eulalia de Nadal, Francesc Posas

## Abstract

Heterogeneity is inherent to living organisms and it determines cell fate and phenotypic variability^1–3^. Indeed, even isogenic cell populations vary in quantifiable phenotypes. Here we generated a high-resolution single-cell yeast transcriptome atlas using genotype and clone RNA barcoded deletions to profile 3500 mutants under control and stress conditions in a genome-scale genetic and environmental perturbation screen. We uncovered a myriad of cell states within a population with specific transcriptional architectures that are both intrinsically and extrinsically regulated, thereby suggesting a continuum of cell states. Cell state occupancy and transition can be genetically modulated with specific mutants that act as state attractors, resulting in differential fitness. By exploiting the power of intra-genetic variability, we identified regulators of transcriptional heterogeneity that are functionally diverse and influenced by the environment.

**One-Sentence Summary:** The yeast single cell-transcriptome atlas based on RNA-traceable genetic perturbations served to map cellular states and define their underlying genetic basis.

## Main

Variability within a population determines cell fate and phenotypic diversity over time (e.g. differentiation, aging)^1^ and facilitates population adaptation to different niches/conditions. Remarkably, even genetically identical cells display variations in growth rate, division, stress resistance, and other quantifiable phenotypes, which lead to global health challenges, such as tumor aggressiveness, chemotherapy resistance, aging, and antibiotic resistance in microorganisms^2,3^. However, this apparent genetic symmetry is not reflected at the transcriptomic or phenotypic level, thereby suggesting that transcription and, the stochastic bursting nature of this process, lead to cell heterogeneity and plasticity^4,5^.

While our ability to profile the transcriptomes of individual cells has increased at unprecedented speed, the molecular mechanisms underlying transcriptional heterogeneity remain a central challenge. Resolving the transcriptome profile of individual cells has been essential in Cell Atlas initiatives seeking mainly to determine the expression programs that identify new cell types and define subtypes/cell states^6^. The concept of cell state has evolved beyond the cell cycle to reflect more dynamic complex cellular behaviors, which are increasingly recognized as being relevant in the transition from health to disease (quiescence, senescence, resistant phenotypes…) ^7–9^. Identifying the molecular basis that defines cell states requires exhaustive cell profiling under both homeostasis and perturbation, an endeavor that will provide the principles to target or engineer specific cell states and, therefore, determine cell population fate.

Genetic screens and extracellular condition profiling have been used to define the relationship between transcriptional heterogeneity and phenotype, mostly using CRISPR knockdown screens in cancer cell lines ^10–13^. Yeast is an ideal reductionist model for assessing transcriptional heterogeneity at the organismal level. In this regard, *Saccharomyces cerevisiae* has pioneered functional screens due to the vast number of molecular tools and genetic resources available for this organism. The amenable yeast genome allows the generation of isogenic genetic perturbations, thereby avoiding the variability generated by CRISPR knockouts or knockdown efficiency using CRISPR inactivation (CRISPRi)^14,15^. One of the most revolutionary tools is the yeast knockout collection (YKOC), in which non-essential genes are deleted from the genome. The YKOC served to dissect genotype-phenotype relationships in bulk assays and has set the groundwork for our understanding of gene network analyses ^16,17^, but has lagged behind in the implementation of single cell transcriptomics^18^.

Here we used yeast as a model organism to define a systematic landscape of gene expression and to understand the principles underlying transcriptional heterogeneity. To this end, we generated a new RNA-barcoded genotype and clone deletion collection and performed a single-cell genome-scale genetic and environmental perturbation screen. This approach allowed us to build a systematic transcriptome atlas, at the genome scale, mapping the complexity of cellular states and functions of the yeast transcriptome and its underlying gene regulatory circuit, as well as defining the genetic drivers of heterogeneity.

## Results

### RNA-traceable deletions enable genome-scale genetic and environmental perturbation screens

To characterize the relationship between genotype and single-cell transcriptomics, we generated a genome-scale library of RNA-traceable deletion mutants by reengineering the yeast knockout collection (YKOC). The YKOC includes deletions of most non-essential genes in the *S. cerevisiae* genome. The structure of the gene deletion cassette to knockout (KO) each gene by homologous recombination consists of a constitutive promoter (p*TEF1*) that drives the expression of *(i)* the *KANMx4* resistance gene and *(ii)* two unique genotype barcodes (20 bp) per deletion; both flanked by common sequences upstream (Uptag) and downstream (Downtag) of the promoter and terminator respectively^16^ (Fig. 1a). This approach renders the genotype identity invisible to transcriptomic readouts and incompatible with single-cell genetic perturbation screens. Inspired by the gene deletion cassette from the original YKOC, we redesigned its structure to generate an RNA-traceable clone and genotype for each mutant (Fig. 1a). Briefly, we generated a PCR cassette to replace the *KAN* resistance marker with *URA3*. This cassette serves two purposes. First, it shortens the heterologous terminator linking the original Downtag barcode to the 3’UTR of *URA3* to a minimum (43 nt) and, second, it adds a clone barcode (5 random nucleotides) downstream of the *URA3* STOP codon (see Methods). This strategy allows the labeling and transcriptional tracing of genotypes and clones (Fig. 1a, Extended Data Fig. 1a). The YKOC was transformed individually, and positive clones were selected (see Methods). The final collection consisted of 4162 mutant strains (82% of the original YKOC) carrying RNA-traceable barcodes suitable for genome-scale genetic screens combined with single-cell transcriptomics (Perturb-seq).

**Fig. 1.**
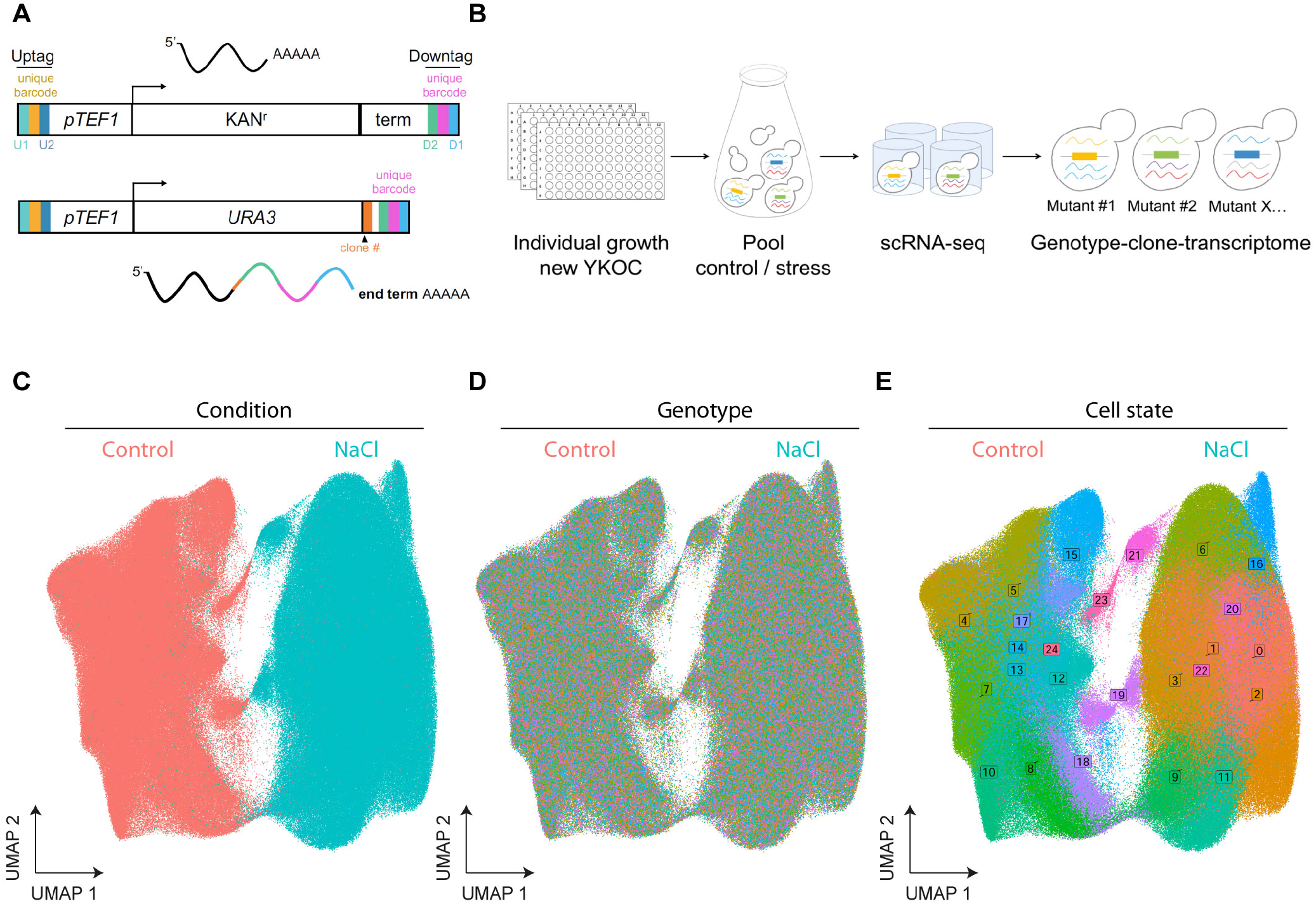
RNA-traceable deletions enable genome-scale genetic and environmental perturbation screens. **a**. Schematic representation of the RNA-traceable yeast knock out collection. Structure of the original deletion cassette (upper panel) and structure of the new RNA-barcoded clone and genotype structure with the resulting position of the barcodes in the 3’UTR. **b**. Perturb-seq experimental layout. Cells were grown individually before being pooled together and subjected or not to stress (0.4M NaCl for 15 minutes). Fixed cells were used for microwell based scRNA-seq. The transcriptome of each cell was linked to the transcriptome through the *URA3* transcript (see methods). **c-e** UMAP of the entire dataset across conditions (c), genotype identify (d) and cell states defined by Seurat (e). UMAPS represent the complete dataset of cels passing the quality chack and assigned to a genotype (n= 710952 cells).

We designed two genome-scale Perturb-seq in which the whole collection of mutants was grown independently (96-well plates) and pooled before being subjected to stress (osmostress; 0.4M NaCl) or control conditions (Fig. 1b). Due to the rapid and transient response of cells to osmostress, we collected and methanol-fixed cells at the peak of the transcriptional response to preserve the transcriptome (see Methods)^19,20^. To generate the scRNA-seq libraries, we used a microwell-based platform for single-cell isolation and oligo dT priming to capture poyladenylated RNA for cDNA synthesis (Yeast GENEXSCOPE HD, Singleron Biotechnologies). We profiled a total of 1.061.865 cells, from which we removed low-quality ones (-/+ 2 standard deviation of mean genes, >10% mitochondrial reads, see Methods). In parallel, for each library, we performed a one-step PCR amplification of the clone and genotype barcode from the *URA3* 3’UTR (Extended Data Fig. 1b). We combined the transcriptome and targeted amplification to assign genotype identity for more than 71% of cells, removing cells with both conflictive and unassigned genotypes from the analysis (Extended Data Fig.1c). The resulting dataset contained more than 3500 mutant genotypes (Extended Data Fig.1d) equally represented across conditions (Extended Data Fig.1e) with an average of 96 and 112 cells/genotype from control and stress conditions, respectively, and had a median coverage of 550 genes/cell and 1200 molecules/cell. Mapping the 3’UTR reads downstream of the genotype barcode allowed us to validate the genomic location of the deleted genes. Overall, 90% of genotype barcodes aligned closely with the endogenous terminator of the deleted gene, thereby confirming genomic loci and thus the robustness of the data (Extended Data Table 1). Correspondingly, we observed a reduced expression of the mutant barcoded gene (Extended Data Fig. 1f). We incorporated clone barcodes into the YKOC to extend its functionality for bulk and single-cell applications. Of note, clones of genotypes with high clonal coverage, the median number of differential expression is 0 (Extended Data Fig. 1g).

To identify sources of intrinsic heterogeneity and avoid normalization biases, we regressed the cell cycle and removed ribosomal genes from the expression matrix (Extended Data Fig.1h). We used highly variable genes as an input to represent the transcriptome of single-cells using Uniform Manifold Approximation and Projection (UMAP) embedding. This approach revealed stress as the predominant clustering factor over genotype identity (Fig. 1c, Extended Data Fig.1i and Fig. 1d). To cluster cells based on their degree of transcriptional similarity, we applied the Louvain algorithm ^21^ to extract the transcriptional signature/gene expression patterns associated with each cell state. Consistent with the predominant effect of stress, the cells in a population were organised into gene expression states within each condition, with few states shared between conditions (Fig. 1e).

Thus, we developed a robust framework that enabled us to build a systematic atlas of the single-cell transcriptome of more than 3500 mutants under control and stress conditions (E-MTAB-14004). Of note, each condition appeared to be determined by a group of cellular states that defines the overall population.

### Cells in a population organize into heterogeneous gene expression states

Given the large-scale dimension of the dataset and the important contribution of cell states, we clustered each condition independently. In the UMAP space, cells distributed into 20 and 18 states in control (0C to 19C) and stress conditions (0N to 17N), respectively (Fig. 2a, 2b). The wild type and 90% of the mutant strains, in both conditions, shared the same distribution across states (Extended Data Fig.2a-d), suggesting that they represent metastable rather than genotype-specific states. We defined the spectrum of transcriptional states by extracting state markers and using the most representative upregulated genes (log2FC >0.25 and *pvalue* <0.05) (Fig. 2c, Extended Data Table 2 and 3). Interestingly, these markers were not markedly distinct, and several genes were shared across states, thereby pointing to a continuum of states within a cell population.

**Fig. 2.**
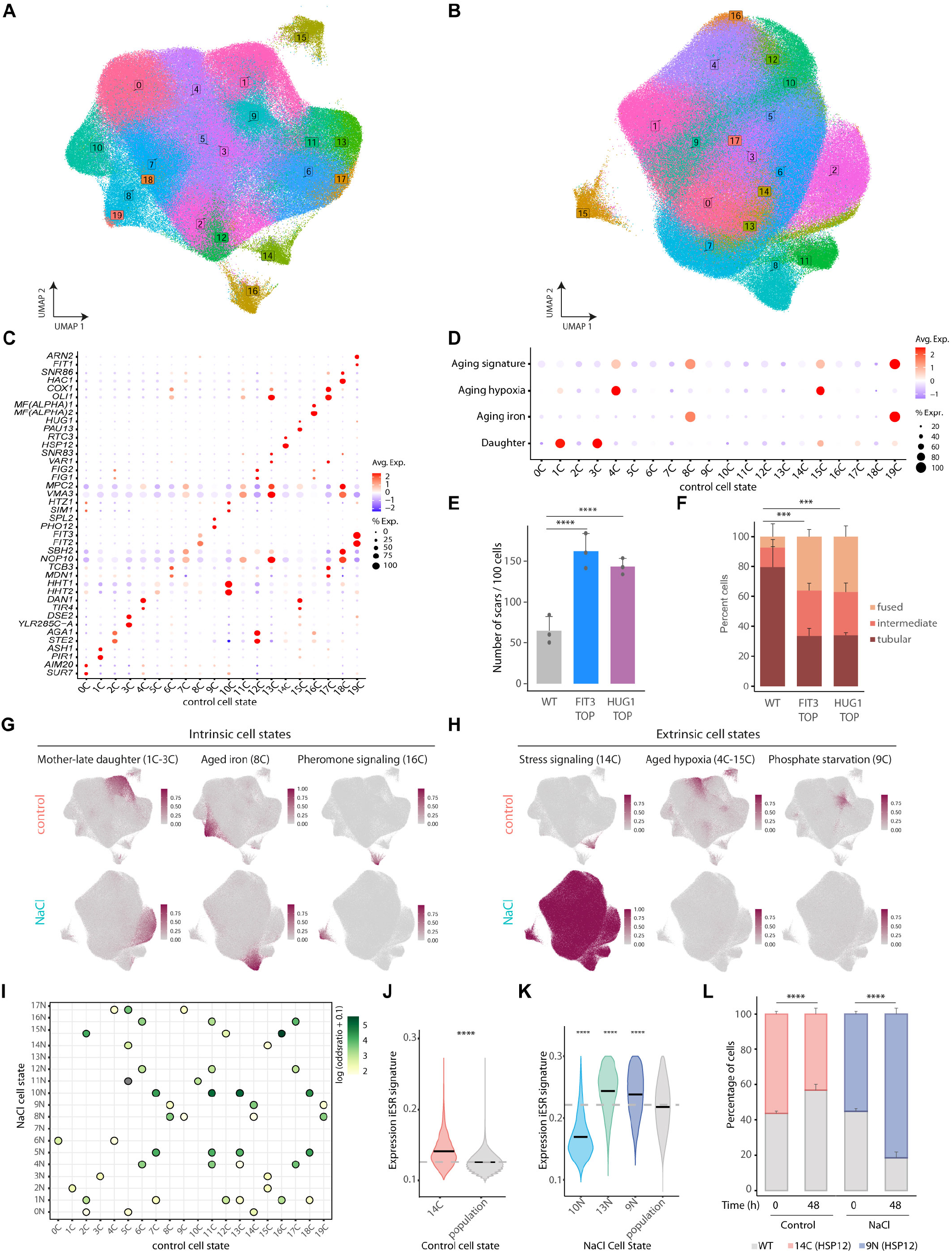
Cells in a population arrange in heterogeneous gene expression states. **a-b**. UMAP of the control (a) and NaCl dataset (b). Cells are colored by cell state indicated by boxed numbers colored according to the corresponding cluster. **c**. Expression of two representative cell state marker genes in the control dataset. Dots size represent the percentage of expressing cells and are colored from high (red) to low (blue) expression. **d**. Expression of the indicated aging signatures across control cell states. **(e)** Number of scars per 100 cells determined by calcofluor staining for a wild type population or the top2% of cells expressing the p*FIT3*-or p*HUG1*-reporters. Data represents the mean of thee independent replicates. Paired t.test is shown agains the wild type (n=3). **f**. Distribution of mitochondrial morphologies for the indicated strains under control conditions by MitoTracker staining (n=3). **g-h**. Examples of intrinsic and extrinsic cell states. UMAP shows the distribution of the cell state signatures for the indicated clusters. Cells are colored by degree of signature expression using two genes per cell state. Darker colors represent higher expression. **i**. Pairwise cell state marker expression correlation across cell states. Per each cell state the expression of upregulated genes was correlated, dots colors represent the degree of similarity determined by Fisher test (darker color indicate higher correlation). **j-k**. Expression of the iESR signature for the indicated cell states compared to the median of population (grey; and grey line). Stars indicate significance comparing against the median. **l**. Competition assay to determine cell state fitness. Cells with the top 2% expression of the *HSP12* reporter were sorted and grown in combination with a wild type cell (labelled with GFP) in rich media (control) or in the presence of stress (NaCl) . Growth of the two populations was assessed at time 0 or after 48 hours by flow cytometry. Data represents mean and standard deviation of three independent experiments and statistical significance is shown respect to the wild type strain (n=3).

In control conditions, cell states involved a variety of functions encompassing the following: cell morphogenesis (e.g., cell wall biogenesis, cell conjugation, pheromone signaling and agglutination genes); metabolic processes (e.g., carbon metabolism, phosphate metabolism, iron metabolism and mitochondrial genes); and protein homeostasis, among others (Extended Data Table 2 and 3). Three distinct clusters were fully separated from the core UMAP and displayed the upregulation of the environmental stress response (ESR) (14C), aging-related signatures (15C), and mating genes (16C) (Fig. 2a). Of note, in addition to 16C, 2C and 12C involve gene signatures of mating pheromone signaling pointing to this pathway as an important contributor to cell state diversity (Extended Data Fig. 2e). Indeed, the heterogeneous expression of pheromone signaling genes, including the detected cell state markers (e.g. *FIG*s, *AGA*s genes), has been found in a similar percentage of the population under normal conditions using nascent transcription reporters, thereby reinforcing the robustness of our dataset^22^.

Among the cell state markers, we recognized well-known transcription programs, such as those that define daughter cells, characterized by the expression of daughter-specific genes (*DSE1-4*, among others^23^. Of note, based on the expression of the *DSE* genes, we distinguished two subpopulations of daughter cells with a differential degree of development; one of them shows the described co-regulation of the *DSE1*,*2* pair with *CTS1* (the chitinase responsible for degrading the mother-daughter barrier) that identifies the most naive daughter cells (3C) versus a second population in which cells expresses the *DSE3*,*4*-genes together with the mother early G1 marker *PIR1*-*HSP150* (1C) (Extended Data Fig. 2e)^24^, thus recapitulating known yeast developmental stages.

Remarkably, clusters 8C and 19C showed a strong upregulation of the iron regulon (*FIT2, FIT3, FET3, ARN1*, among others), mirroring the hallmark signature of aged cells defined by bulk RNA-seq ^25^ (Fig. 2d) and correspondingly, accumulated iron^25^. Of note, while 8C highly expressed the iron regulon, 19C encompassed a smaller number of cells and a larger number of genes involved in cell wall organization induced during stress conditions, perhaps representing a later aged stage and thus indicating that the data allows the tracing of different degrees of phenotypical development. Moreover, extending the aging signature defined in bulk to the top 10 genes and distinguishing between iron-related and unrelated genes, clusters 4C and 15C also stood out, revealing a bimodality within aging states in the cell population. This second module (4C and 15C) included the upregulation of genes known to be induced under hypoxia (*TIR, DAN* and *PAU* families) but not the upregulation of iron genes (Fig. 2d and Extended Data Fig. 2f). Thus, scRNA analysis suggests divergent paths towards aging through iron metabolism or hypoxia. Correspondingly, cells expressing either of the differential aging reporter genes (i.e. *FIT3* or *HUG1*) showed a similar aging phenotype, for instance, an increased number of scars (Fig. 2e) and fragmented mitochondrial morphology (Fig. 2f). Taken together, our data suggest that, within a cell population, coordinated heterogeneous gene expression patterns lead to interconnected cell states which can generate distinct cell fates.

### Transcriptome mapping unveils intrinsically and extrinsically metastable cell states

To assess whether cell states are conserved in different conditions, we assessed the degree of transcriptional correlation across cell state markers across conditions. We classified “intrinsic states” as those whose gene expression patterns were reciprocally represented in both conditions (e.g., daughter, aged cells), while “extrinsic states” were defined as those differentially modulated depending on the condition (i.e. stress)(Fig. 2g and 2h). A large proportion of states in the control condition (15 out of the 20) had a counterpart upon exposure to stress (Fig. 2i). Intrinsic states included developmental and morphogenic programs, the aging states, and states related to mitochondrial function. These observations thus indicate a high degree of conservation of the core clusters across conditions.

Extrinsic states arose either because the expression of cell state markers are modulated upon stress losing their specificity, or because novel stress-regulated states appear during adaptation. For example, under stress, markers of the hypoxic aging and phosphate metabolism clusters (4C-15C and 9C, respectively) were widely expressed, suggesting altered cell wall composition and metabolism (Fig. 2h). On the other hand, some stress-dependent cell states showed a weak correlation with control states due to the expression of stress-related markers genes. These involved multiple functions, including carbon metabolism (clusters 0N and 13N), ADP metabolism (cluster 14N), and protein folding (cluster 17N), thereby suggesting that clusters that arise in the presence of stress are related to specialized protective functions (Fig. 2i, S2g and Extended Data Tables 2 and 3). For example, the stress cluster 9N correlated with the basal-stress cluster in the control condition (14C) and both displayed higher expression of the induced Environmental Stress Response (iESR) (e.g *HSP12*) identified by bulk RNA-seq ^26,27^ (Fig. 2j, 2k and Extended Data Fig. 2h). The expression of the stress response programme (14C) resulted in a decreased fitness compared to wild type under normal conditions (Fig. 2l), however resulted in a fitness advantage in the presence of stress (Fig. 2l). Of note, cluster 13N, which was also hyperesponsive, was enriched in daughter cells (Extended Data Fig. 2l). A global comparison of the response of mother and daughter cells revealed that the latter systematically showed a stronger transcriptional response to stress (Extended Data Fig. 2i), thereby suggesting that cell states determine the plasticity of the cell to environmental conditions.

Therefore, the Yeast Tanscriptome Atlas identifies a set of interconnected cell states within an isogenic population that reflects physiology and function. These cell states represent an essential trait (intrinsic) that is robust to genetic perturbations but dynamic to environmental perturbation (extrinsic).

### Cell state occupancy can be genetically regulated

The transcriptional signatures underlying cell states provide genotype-phenotype information. To understand the genetic determinants of cell states, we assessed the enrichment or depletion of specific genotypes over the defined cell states (Extended Data Tables 4 and 5). The wild type and most mutants (approx. 90%) were not biased towards any cell state in either condition tested (Fig. 3a-3b and Extended Data Fig. 3a-3a) whereas 10% of mutants were significantly enriched in or depleted of specific cell states (253 in control and 331 in stress; with 162 of them common to both conditions) (odds ratio >1, *pvalue* < 0.05) (Extended Data Fig. 3c, 3d). Cell-state biased mutants tended to be enriched in or depleted of a single state and rarely in multiple states. Although accumulation towards particular states can be either plastic (condition-dependent) or genetically determined (condition-independent) and it is clone-independent (Extended Data Fig. 3e, 3f). Our results suggest that cell state organization is a fundamental feature that is resistant to most genetic or environmental perturbations.

**Fig. 3.**
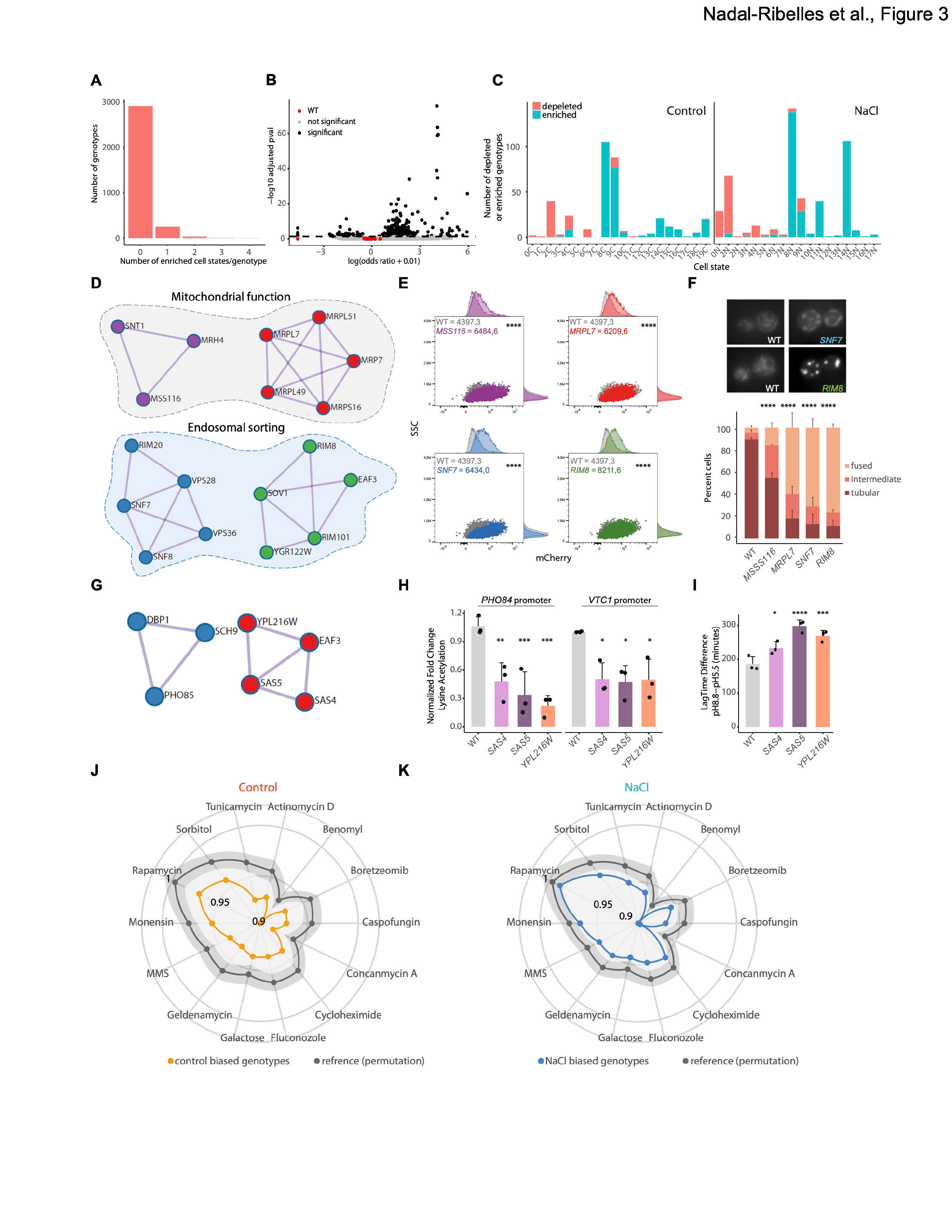
Transcriptome mapping unveils intrinsically and extrinsically metastable cell states. **a**. Distribution of the number of enriched cell states per genotype in control conditions. **b**. Volcano plot shows the occupancy of each genotype (mutants in black and wild type clones in red). Black line and dots show the threshold for statistical significance (adjusted *pvalue* 0.05) in control conditions. **c**. Stacked barplot shows the number of genotypes statistically enriched (blue) or depleted (red) per each cell state in each condition. **d**. Protein enrichment network of genotypes in control condition enriched in cluster 8C (odds ratio >1 and adjusted *pvalue* <0.05). Nodes are colored according to module enrichment (MDCE, see Methods). **e**. Expression of the p*FIT3* reporter (cluster 8C) in the selected mutants assessed by flow cytometry, expression of the mCherry reporter is shown in the x axis agains the SSC for the indicated strains colored as in (d). Histograms represent the distribution of the scatter plot axis of a representative experiment of three independent experiments and the mean, standard deviation and significance to the wild type (paired t.test) is shown within (n=3). **f**. Representative images of mitochondrial morphology stained by MitoTracker for the indicated strains. Stacked barplots show the distribution of mitochondrial morphology under control conditions for the selected mutants. Graphs represent a the distribution and standard deviation of 150 cells from each strain (50 cells per independent replicate, n=3). **g**. Protein protein interaction for genotypes enriched in cell state 9C. Nodes are colored according o their MDCE. **h**. Levels of total Lysine acetylation were determined by ChIP at the indicated strains in control conditions for the indicated promoters. Data is represented normalized as Fold Change respect to the wild type which is set to 1. Bars represent the mean and standard deviation of three independent biological replicates (dots). Paired t.test is shown respect to the wild type. **i**. Growth of the indicated strains was assessing growth curves in control (CSM pH=5.5) or alkalyine conditions (pH=8.8). The bars represent the difference in Lag time of each strain between the alkalyine media respect to control media of three independent experiments (n=3). Paired t.test is shown respect to the wild type. **j**. Radar plot shows the fitness score across a stressor panel for mutants enriched in specific cell states in control conditions (orange) against a random permutation as a reference (grey). Light grey ribbons show the 95% interval confidence. **k**. Fitness score as in J for mutants enriched in cell states in stress conditions (blue).

We then analyzed the distribution of the biased mutants across cell states. Two related states were observed to be preferentially enriched by mutants in control and stress conditions (8C and 9C in control and 8N and 14N in stress) (Fig. 3c and Extended Data Fig. 3g-3h). Indeed, cluster 8C (related to 8N) displayed the aged iron-regulon signature, and mutants that impair mitochondrial functions (mitochondrial translation and ATP transport) and iron homeostasis accumulated in these clusters and displayed this iron-dependent premature aged signature in both conditions (Fig. 3d). These findings suggest, and reinforce the initial observations, that defects in mitochondrial function lead to impaired iron homeostasis^28^ leading to a premature aged transcriptome. Of note, mutants defective for endocytosis (i.e., endosomal sorting and ESCRT complexes) also accumulated in the same state (Fig. 3d). We verified the occurrence of cell state bias by comparing the percentage of iron reporter-expressing cells (p*FIT3* mCherry) in the mitochondrial and endocytosis-related mutants in comparison to the wild type. Indeed, these mutants showed higher expression of the *pFIT3* reporter (Fig. 3e) and displayed an aging phenotype (i.e., fragmented mitochondria) (Figure 3F), thereby suggesting that such phenotypes can be promoted by alterations in gene function. This results are in agreement with the newly identified role of Snf7, a member of the ESCRT complex, in macromitophagy^29^. Thus, our data highlights the potential of using transcriptional phenotypes to define novel phenotype-genotype interactions.

The transcriptional response to genetic perturbations also provides information-rich profiles to naturally infer and predict gene function. For example, in the control condition, the second cell state with the larger number of biased mutants (9C) showed a strong transcriptional signature related to phosphate starvation (Figure 2h)^30^. Accordingly, genotypes depleted of this cluster include *hmt1*, a mutant in an arginine methyltransferase and known positive regulator of phosphate genes^31^, whereas genotypes that accumulated in this cluster include known negative regulators of phosphate metabolism (e.g *PHO85, KCS1*)^32,33^ (Fig. 3g and Extended Data Fig. 3e). Additionally, we identified other mutants involved in histone acetylation, namely the SAS (Sas4, Sas5 a trimeric HAT complex) and Nua4 (*EAF3*) complexes, with a increased expression of the phosphate genes, suggesting a novel role for these complexes in phosphate homeostasis (Fig. 3g). Within the protein interaction network that contained the SAS and NuA4 complexes, we identified *YPL216W*, a paralog of unknown function of *ITC1*, a component of the Isw2 chromatin remodeling complex involved in gene silencing. Correspondingly, the deletion of *YPL216W* leads to a decrease in lysine acetylation at promoters of phosphate genes (i.e., *PHO84* and *VTC1*)(Fig. 3h) and result in similar phenotypes to that observed in SAS mutants, for example slower growth in alkalyine media (Fig. 3i), validating a novel set of downstream targets for these complexes.

To determine the overall phenotypic effects of cell state confinement, we evaluated the fitness of all mutants with a biased cell state distribution (253 from control and 331 from stress) using available environmental screens across 14 stressors^34^. Cell state-biased mutants from both conditions systematically displayed lower fitness scores compared to a random permutation, regardless of the stressor type (Fig. 3j-3k). These observations suggest that transcriptional homeostasis favors cell state organization enabling correct fitness.

Therefore, by integrating single-cell transcriptome data, cellular state mapping, and genetic perturbations, we have defined the regulatory logic that governs cell states and defined several specific mutants that act as state attractors. The Yeast Transcriptome Atlas provides a unique resource that links genotype-transcriptome-phenotype, offering new insights into gene function that are difficult to discern at the population level.

### Different cellular functions drive transcriptional heterogeneity under control and stress conditions

Transcriptional heterogeneity is a source of cell plasticity and it has an impact on cell phenotype. Leveraging on the single-cell resolution of the Perturb-seq, we reasoned that scoring the degree of transcriptional heterogeneity shown in each mutant would reveal genetic drivers of heterogeneity and their conservation across conditions. By applying an SVD-based leverage score ^35^, we determined the genetic perturbations that resulted in significantly deviated gene expression from the wild type. Of note, the average leverage score strongly correlated across conditions (Extended Data Fig. 4a), and is independent of cell number (Fig. S4b-S4c). We then used the standard deviation of the scaled leverage score to identify mutants with increased (negative regulators) or decreased (positive regulators) heterogeneity, using a 30% threshold relative to the wild type (see Methods). While most mutants, in both conditions, did not lead to changes in transcriptional heterogeneity, we identified a larger fraction of negative regulators of transcriptional heterogeneity (approx. 150 mutants) than positive regulators (approx. 20 mutants) (Fig. 4a-4b) (Data S6). This pattern mirrors observations in the human Perturb-seq datasets ^35^(Extended Data Fig. 4d and 4e), thereby suggesting that transcriptional heterogeneity is kept within a defined range and only a few mutations alter it.

**Fig. 4.**
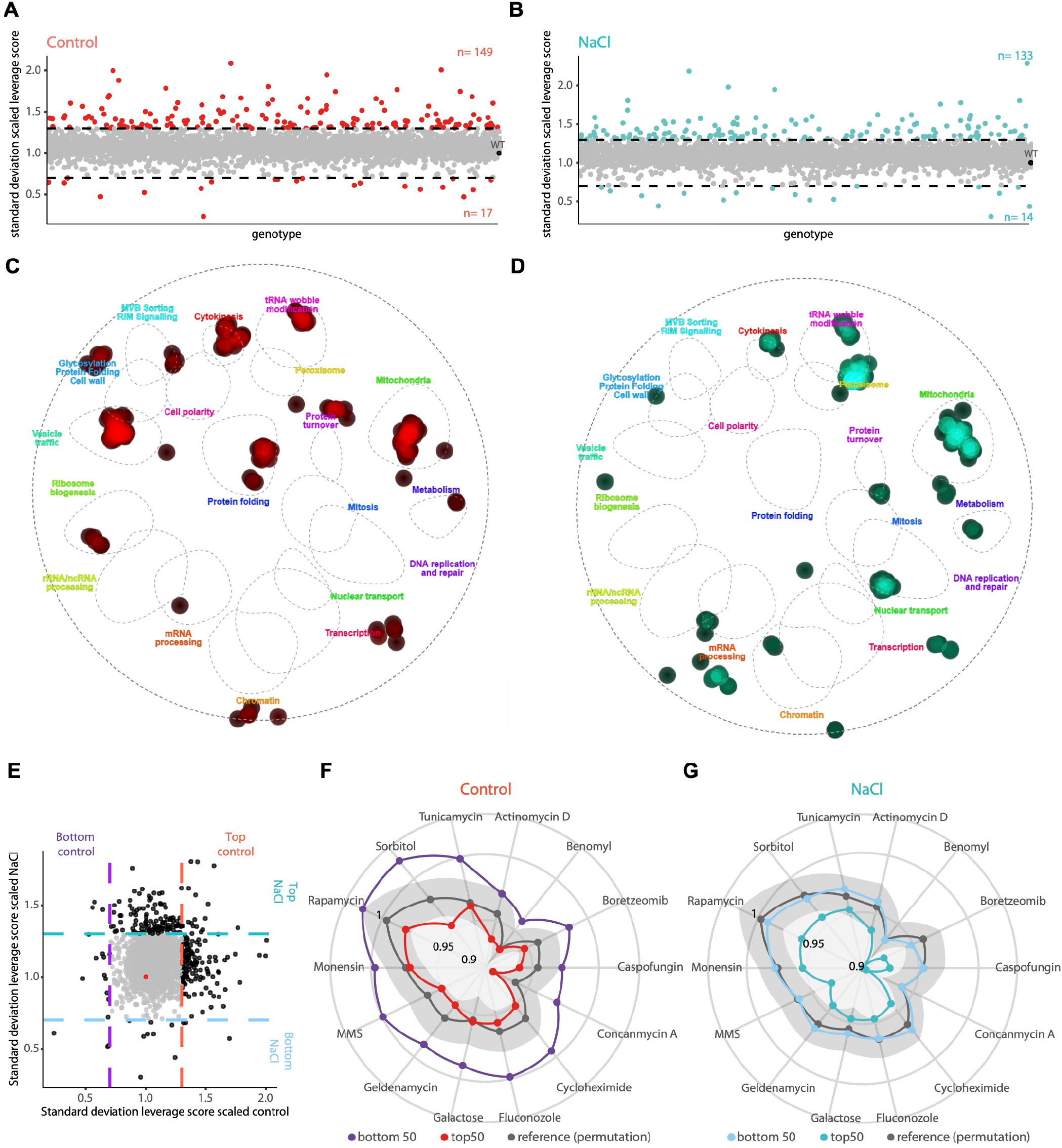
Cell state occupancy can be genetically regulated. **a-b**. Distribution of the standard deviation of heterogeneity (scaled leverage score, y axis) per each genotype (x axis) in control (a) and stress (b). Dashed lines indicate the >30% increase or decrease threshold compared to the wild type. Colored points represent positive or negative regulators. The total number of genotypes above and below threshold is shown in graph. **c-d**. Projection of negative regulators of heterogeneity per each condition (control, c and NaCl, d). The list of candidates was overlayed with the yeast genetic interaction network to associate genes to function. Points represent the density of mutants over each indicated cellular function from the cell map. **e**. Comparison of transcriptional heterogeneity per genotype (standard deviation scaled leverage score). Each point represents a genotype in control (x axis) or stress (y axis). Points are colored according to the >30% difference respect to the wild type. **f-g**. Radar plot shows the fitness score across a stressor panel for mutants of top50 (red and turquoise) and bottom 50 (purple and blue) genotypes with increased/decreased heterogeneity per each condition (f, control and g, NaCl) against a random permutation as a reference (grey). Light grey ribbons show the 95% interval confidence.

To understand the nature of the potential drivers of heterogeneity, we projected the negative regulators of each condition individually onto the yeast genetic interaction map^36,37^. Negative drivers spanned a variety of functions in both conditions. As expected, under control conditions, chromatin, transcription, and translation mutants were present, but surprisingly, genes related to vesicle trafficking, cytokinesis, mitochondrial function, and tRNA modifications were also included (Fig. 4c). Of note, mutants related to mitochondrial dysfunction generate increased heterogeneity in mammals^35^. Upon stress, we found that negative regulators involved some specific cellular functions, such as peroxisome function, nuclear transport and chromatin/transport. However, mutants related to mitochondrial function were also among the most represented drivers (Fig. 4d). Of note, while both control and stress drivers showed shared functions, such as mitochondrial function, the identity of these genes poorly overlapped (10%, Fig. 4e and Extended Data Fig. 4f and 4g). Therefore, transcriptional heterogeneity is regulated by several core processes, and it is condition-specific, suggesting that the identification of molecular drivers should be performed for each specific condition of study.

To assess the physiological impact of transcriptional heterogeneity, we extracted cell fitness scores from the top and bottom 50 drivers for each condition from 14 phenotypic screens, as shown in Fig. 3j-3k^34^. Mutants with decreased heterogeneity under control conditions showed significantly higher fitness upon stress than genotypes with high variability^38^ (Fig. 4f). Conversely, mutants with high transcriptional heterogeneity in osmostress displayed consistently lower fitness in most stress conditions (Fig. 4g). These observations suggest that transcriptional heterogeneity is a quantitative trait that influences cell fitness. Our results support the notion that excessive transcriptional heterogeneity generates deviated transcriptional patterns from the wild type that ultimately render cells vulnerable to stressors, thereby weakening cell adaptability.

## Discussion

Here we performed a single-cell genome-scale Perturb-seq in *S. cerevisae* by modifying the initial structure of the (non-essential) YKOC to generate RNA barcoded mutants, which allowed us to exploit the genotype-transcriptional phenotype, including clonal resolution. This approach enabled us to generate a genome-scale single-cell Yeast Transcriptome Atlas. The resulting dataset is consistent with and significantly extends the previous large-scale transcriptomic profiling in yeast, which covered approximately 25% of the yeast deletome in bulk under control conditions ^39^. The atlas served to identify ubiquitously shared transcriptional states within a cell population. These, were present in more than 90% of the mutants analyzed, thereby indicating the strong robustness of cell states against genetic perturbations.

We categorized cell states into “extrinsic”, which are modulated by external conditions and promote specialized adaptive functions, and “intrinsic”, which are genotype- and condition-independent. These transcriptional states involve developmental and core functional transcriptional programs (oxidative phosphorylation, cell cycle, cell wall morphogenesis, protein homeostasis, and stress responses, among others). Interestingly, some of the intrinsic states associated with protein homeostasis, cell cycle, oxidative phosphorylation and stress response are also found in human Perturb-seq ^35^, thus indicating partial conservation of functional cell states. The analyses of the transcriptional programs in each state served to deconvolute different degrees of phenotypical development or distinct transcriptional paths that lead to aging phenotypes.

The high amount of information provided by this Perturb-seq yields a comprehensive snapshot of cellular states as a function of genotype and condition. We harnessed the potential of 10% of the mutants that showed biased cell state accumulation. Some mutations irreversibly confined cells to specific transcriptional states, acting as stage attractors, whilst others, depending on the environmental conditions, relocated or pushed cells into specific states. The transcriptional response of the mutants also provides information to infer and predict gene networks and functions. For example, in the control condition, the second cell state with the larger number of biased mutants displayed a strong transcriptional signature related to phosphate metabolism and this served to identify novel regulatory layers of phosphate homeostasis.

In addition to uncovering the genetic underpinning of cell states, the catalog of transcriptional phenotypes captures genotype-specific gene expression patterns, thus enabling the identification of positive and negative regulators of transcriptional heterogeneity. Whereas in higher eukaryotes these have been linked mainly to changes in copy number, our data in yeast suggest that negative regulators involve several cellular functions, including transcription, translation and metabolic functions. Of note, some of the most abundant negative regulators involved several mitochondrial functions. Similarly, perturbation of mitochondrial dysfunction in the human Perturb-seq leads to increased heterogeneity ^35^. These results underscore the importance of considering both global and condition-specific effects when assessing the impact and regulation of transcriptional heterogeneity. In summary, the Yeast Transcriptome Atlas provides a reference map, enabling both hypothesis-driven and hypothesis-generating exploration of cellular behaviors at several levels, such as genotype-phenotype relationships, and it can be used to identify conserved traits between eukaryotes.

## Acknowledgments

We thank the partnership with Singleron Singleron Biotechnologies GmbH for generously lending the Matrix equipment, the Singleron Bioinformatic service (Jaren Sia and Stacy Xu) for analysis and the Singleron R&D (Julie Laliberte) for technical support. We also would like to thank Dr. Cayetano González (IRB Barcelona), the IRB Functional Genomics Core Facility, the Flow Cytometry Unit from the Scientific and Technological centers (CCiTUB) and staff members Jaume Comas and Ricard Alvarez, as well as Oscar Reina and Lidia Mateo (Biostatistics Unit IRB Barcelona) for technical support. We also thank Sandra Clauder and Dr. Cosimo Jan (from Dr. Steinmetz group) for generously sharing the original Yeast Knock Out collection. Additionally, we thank Aitor González, Dr. Pablo Latorre, Dr. Holger Heyn and Dr. Ivo Gut for helpful discussions during the initial conceptualization of the project. We would like to thank the Associación Española Contra el Cancer for supporting ADV through the AECC Excelencia program. Finally, we would like to thank Aida Fernández for technical support and special thanks to Mònica Romo for instrumental assistance in conducting the experimental validations.

## Author contributions

Conceptualization: MNR, CS, LS, EdN, FP

Investigation: MNR, CS, AD, CS-O, YM

Funding acquisition: MNR, EdN, FP

Supervision: EdN, FP

Writing – review & editing: MNR, CS-O, LS, EdN, FP

## Data availability

Raw and processed scRNA-seq is composed of transcriptome data (Singleron libraries), targeted amplification are deposited at Array Express. Additionally, and fully preprocessed samples as Seurat object for the complete dataset combined or split objects are available through Synapse. All the code generated for the analysis and figures can be accessed at the Yeast Transcriptome Atlas repository in github that contains the code to reproduce the figures and tables generated in the figures including those already provided as supplementary data;

