## supplemental figures and methods for "Perturbation-driven transcriptional heterogeneity impacts cell fitness"

**The file includes:**

Extended Data Figures 1 to 4
Materials and Methods
Tables 1 and 2
  Table 1: Plasmids generated in this study
  Table 2: Yeast strains generated in this study
Methods References

**Extended Data (for this manuscript include the following:**

Extended Data Table 1: Distance to expected chromosome location of each genotype
Extended Data Table 2: Table of cell state defining genes in control.
Extended Data Table 3: Table of cell state defining genes in NaCl.
Extended Data Table 4: Table cell state enrichment per genotype in control.
Extended Data Table 5: Table of cell state enrichment per genotype in NaCl.
Extended Data Table 6: Leverage score metrics per each genotype in both conditions.
Extended Data Table 7: List of genes used to score gene signatures.

Nadal-Ribelles et al., Extanded Data Figure 1

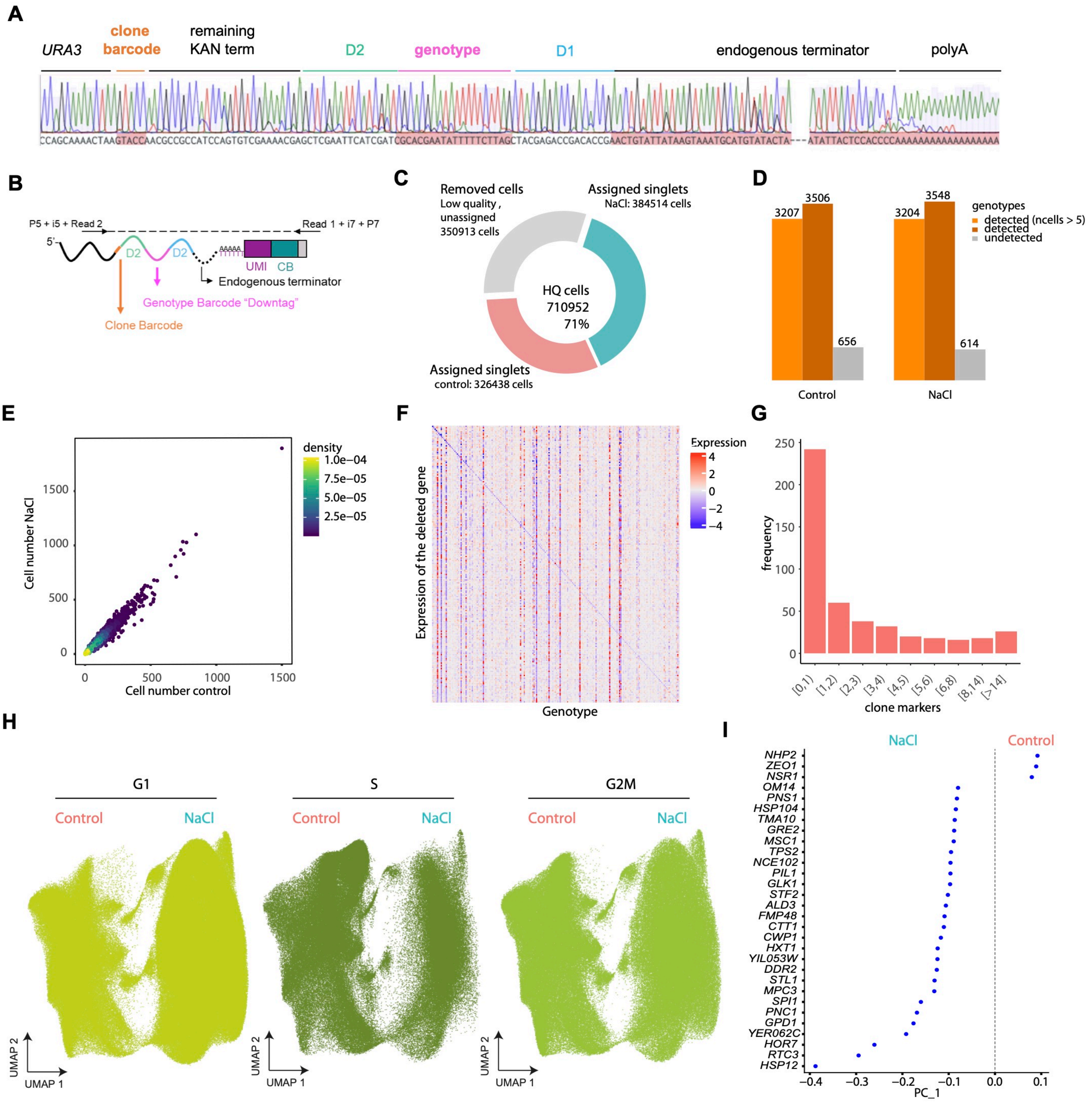

**Extended Data Fig. 1. RNA-traceable deletions enable genome-scale genetic and environmental perturbation screens.**

**a.** Schematic representation of the RNA-barcoded structure and a representative Sanger sequencing chromatogram. **b.** Overview of the targeted amplification PCR strategy, the *URA3* transcript and the contained barcodes are shown in colors. Rectangle represents the anchoring oligo dT used for cDNA synthesis. Arrows indicate the location of primers and dotted line the amplification product. **c.** Distribution of cells in the entire dataset after quality check (QC). Plot represents a total of 1.061.865 cells. Colored regions show removed cells (low quality cells or doublet cells, grey) or cells that pass the QC based per each condition. The total number per each section and kept for further analysis are shown. **d.** Genotype coverage per each condition is shown based for all detected genotypes (dark orange), all detected genotypes with >5 cells used for the analysis (light orange) and unassigned cells based on the 4162 possible genotypes. **e.** Correlation of cell number per genotype across conditions. Points are colored by density, warmer colors indicate higher density. **f.** Heatmap representing the expression of a deleted gene (y axis) against the inferred genotype (x axis) (cold colors indicate low expression and warmer higher). Top 200 expressing genotypes are shown. **g.** Distribution of the number of differentially expressed genes within clones for 220 genotypes with more than 200 cells 9 clone in control conditions (see Methods). **h.** UMAP of the entire dataset colored by the indicated cell cycle phase. Cell cycle was determined using Seurat and canonical phase specific genes (see Methods). **i.** Gene loadings of dimensionality reduction for Figure 1F, green shade indicates loading for the stress cluster and red genes loading control cluster for PC-1.

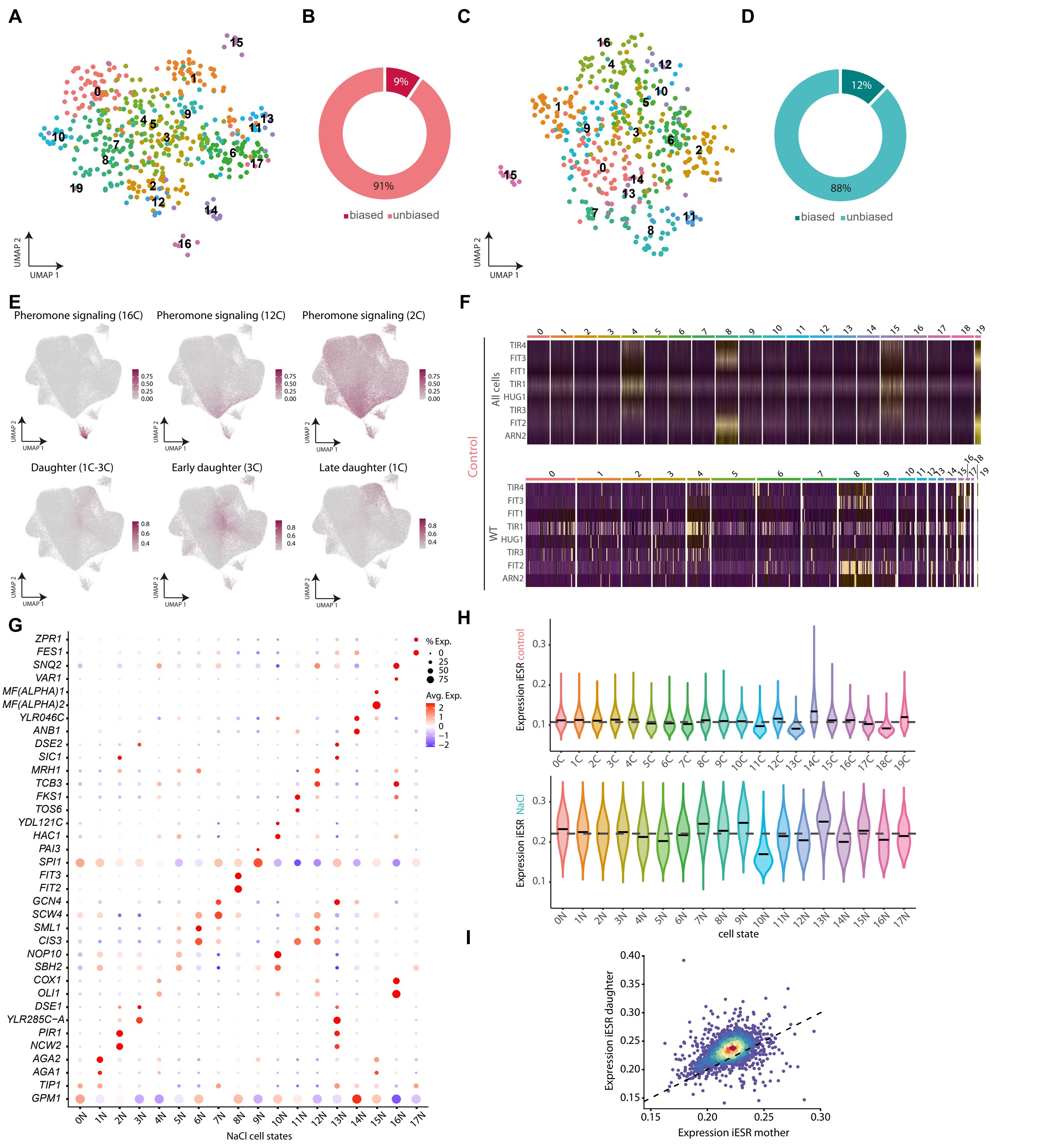
A
B
9%
91%
biased unbiased
C
D
12%
88%
biased unbiased
UMAP 1
UMAP 2
E
Pheromone signaling (16C)
Pheromone signaling (12C)
Pheromone signaling (2C)
Daughter (1C-3C)
Early daughter (3C)
Late daughter (1C)
F
Control
All cells
WT
TIR4
FIT3
FIT1
TIR1
HUG1
TIR3
FIT2
ARN2
G
ZPR1
FES1
SNQ2
VAR1
MF(ALPHA)1
MF(ALPHA)2
YLR046C
ANB1
DSE2
SIC1
MRH1
TCB3
FKS1
TOS6
YDL121C
HAC1
PAI3
SPI1
FIT3
FIT2
GCN4
SCW4
SML1
CIS3
NOP10
SBH2
COX1
OLI1
DSE1
YLR285C−A
PIR1
NCW2
AGA2
AGA1
TIP1
GPM1
% Exp.
Avg. Exp.
NaCl cell states
H
Expression iESR control
Expression iESR NaCl
cell state
I
Expression iESR daughter
Expression iESR mother

**Extended Data Fig. 2. Cells in a population arrange in heterogeneous gene expression states.** **a.** UMAP of wild type cells extracted from the control dataset colored and labeled by cell state. **b.** Percentage of mutants in control condition with unbiased cell distribution (light red) or biased distribution (dark red). **c.** UMAP of wild type cells as in (a). **d.** Percentage of mutants according to their distribution unbiased (light blue) or biased (dark blue). **e.** UMAP representing the expression signature defined by two genes of control clusters related to pheromone signaling (clusters 16C, 12C and 2C, upper panel). Projection of daughter cell state markers clusters (1C and 3C independently and combined). Darker color indicates higher expression. **f.** Heatmap represents the expression of the aging signature for a downsampled subset of the entire control dataset (upper panel) and wild type cells only (lower panel). g. Expression of two representative cell state marker genes of the stress dataset. Dot size represent the percentage of expressing cells and are colored from high (red) to low (blue) expression. **h.** Expression of the iESR signature for all cell states in each condition compared to the median of population (grey line). **i.** Scatter plot shows the expression of the iESR signature (n=175 genes) per each genotype as a function of generation (mother cells, x axis and daughter cells y axis). Each point represents a genotype and points are colored by density (warmer colors indicate higher density).

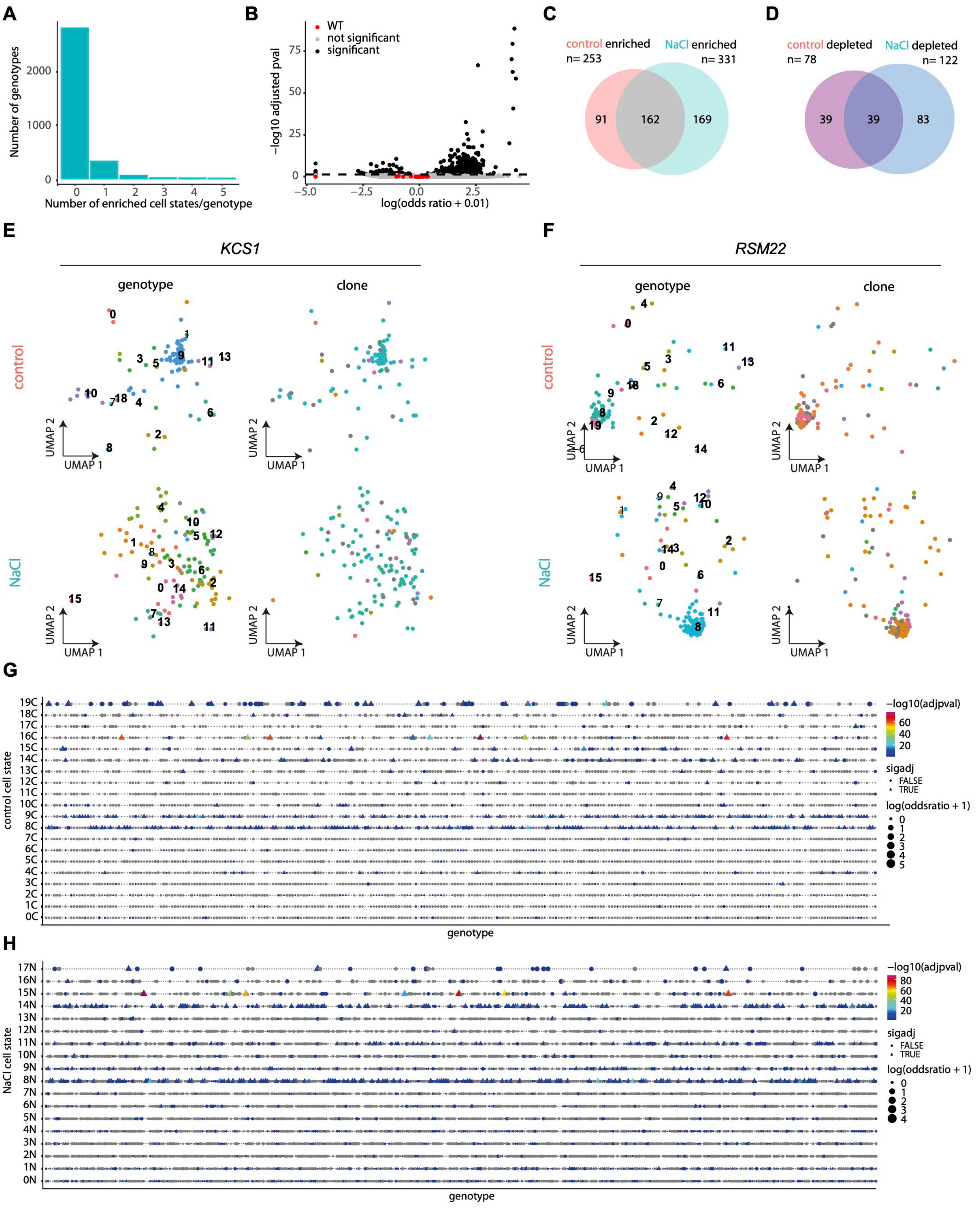
A
Number of genotypes
Number of enriched cell states/genotype
B
WT
not significant
significant
−log10 adjusted pval
log(odds ratio + 0.01)
C
control enriched
n= 253
NaCl enriched
n= 331
91
162
169
D
control depleted
n= 78
NaCl depleted
n= 122
39
39
83
E
KCS1
F
RSM22
genotype
clone
control
NaCl
UMAP 1
UMAP 2
G
control cell state
genotype
−log10(adjpval)
sigadj
FALSE
TRUE
log(oddsratio + 1)
H
NaCl cell state
genotype
−log10(adjpval)
sigadj
FALSE
TRUE
log(oddsratio + 1)

**Extended Data Fig. 3. Transcriptome mapping unveils intrinsically and extrinsically metastable cell states.**

**a.** Distribution of the number of significant cell states per genotype in stress conditions. **b.** Volcano plot shows the occupancy of each genotype (mutants in black and wild type in red). Black line and dots show the threshold for statistical significance (adjusted *pvalue* 0.05) in control conditions. **c-d.** Overlap between cell state enriched (c) or depleted (d) genotypes across conditions (adjusted *pvalue* <0.05). **e-f.** UMAP representation of genetically determined cell state for the indicated mutants across conditions (top/bottom panels). Points are colored and labeled based on the Seurat cluster (left panel) or based on their clone identity (grey points represent unassigned clones). **g-h.** Phenomap shows the top500 genotypes with biased cell state enrichment (x axis) as a function of cell state (y axis) in control (upper panel) and stress (lower panel). Shape indicates statistical significance and are colored according to the odds ratio.

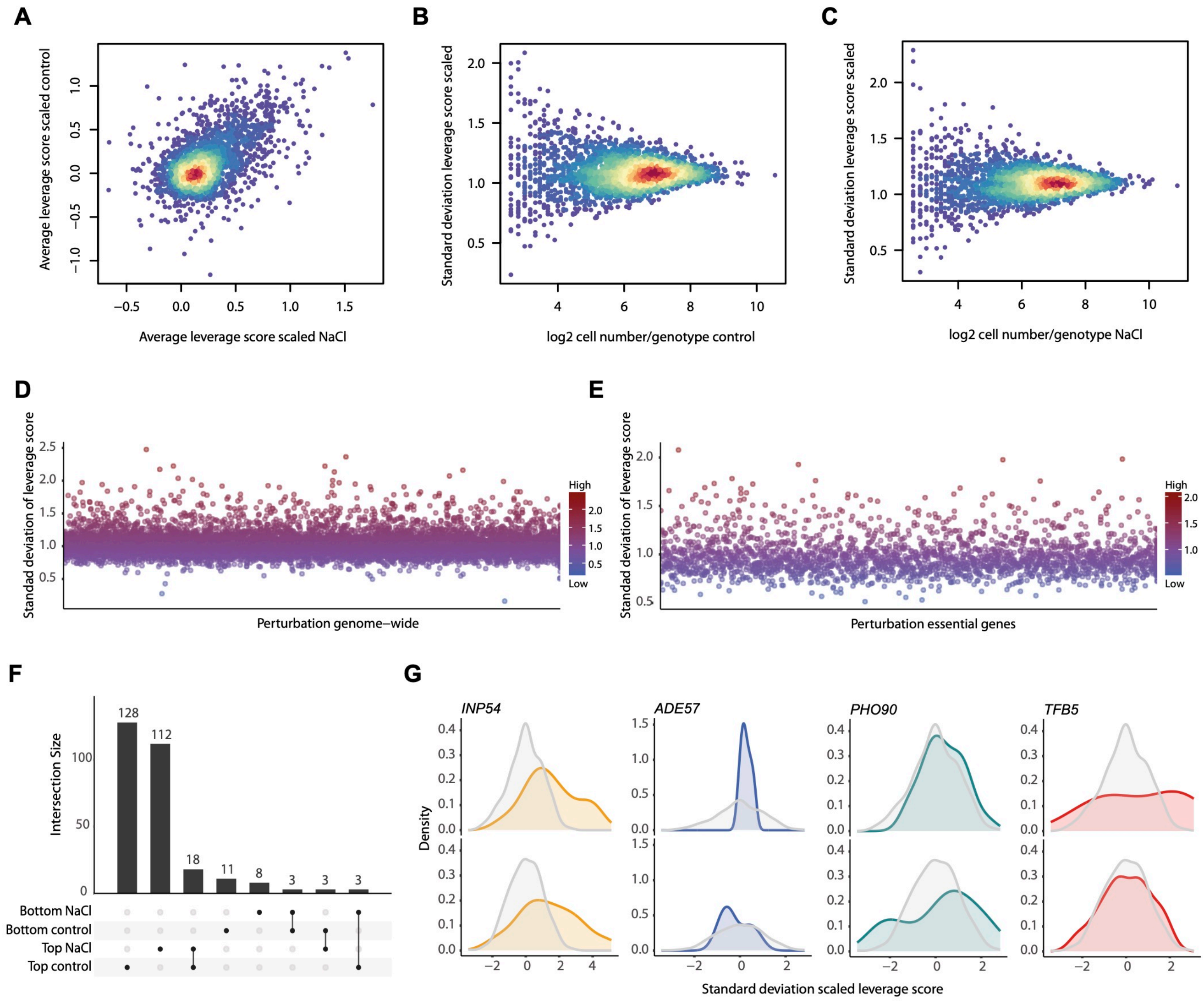
A
Average leverage score scaled control
Average leverage score scaled NaCl
B
Standard deviation leverage score scaled
log2 cell number/genotype control
C
Standard deviation leverage score scaled
log2 cell number/genotype NaCl
D
Standad deviation of leverage score
Perturbation genome−wide
High
Low
E
Standad deviation of leverage score
Perturbation essential genes
High
Low
F
Intersection Size
128
112
18
11
8
3
3
3
Bottom NaCl
Bottom control
Top NaCl
Top control
G
INP54
ADE57
PHO90
TFB5
Density
Standard deviation scaled leverage score

**Extended Data Fig. 4. Cell state occupancy can be genetically regulated.**
**a.** Correlation of the average leverage score per each genotype (blue dots) in control (x axis) and stress (y axis) conditions. **b-c.** Distribution of standard deviation of scaled leverage score (y axis) against the number of cells per genotype in control (b) and NaCl (c) datasets. Point density is shown in warmer colors (high) and colder colors (low). **d-e.** Distribution of the standard deviation of scaled leverage score (y axis) in human Perturb-seq in K562 genome-scale (d) and essential gene screens (e). **f.** Upset plot shows the overlap between negative and positive regulators identified in each condition. Bars are ordered in descending order in each indicated category and the total number is shown above. **g.** Representative distributions of transcriptional heterogeneity (leverage score) patterns for the indicated mutants (colored distributions). The background filled distribution in grey is shown for the wild type strain in each conditions.

### Methods

#### Generation of a single cell transcription atlas by using a modified yeast knock out collection.

We generated a genome-scale library of RNA-traceable deletion mutants by reengineering the yeast knockout collection (YKOC)[1]. Briefly, we generated a PCR cassette to replace the G418 resistance marker with URA3 to shorten the heterologous terminator linking the original Downtag barcode to the 3'UTR of *URA3* to a minimum (43 nt) and added a clone barcode (5 random nucleotides) downstream of the *URA3* STOP codon (see below). This strategy allows the labeling and transcriptional tracing of genotypes and clones.

#### Oligonucleotides used for the modification of the YKOC collection.

Oligonucleotides for strain generation (Integrated DNA Technologies) were purified by PAGE purification and resuspended with nuclease free water (Thermo, 10977035) at a final concentration of 100 µM.

*For YKOC deletion strains:*

*OMN761:*
CACATCACATCCGAACATAAACAACCatgggtaaggaatcgaaagctacatataaggaac
*OMN773:*
TCGATGAATTCGAGCTCGTTTTCGACACTGGATGGCGGCGTTNNNNNttagttttgctggccgc
atc
*Primers to generate WT strains:*
*OMN774:*
GATTCGGTAATCTCCGAGCAGAAGGAAGAACGAAGGAAGGAGCAGACATGGAGGC
CCAGAATACC
*OMN775:*
ATTTGTGAGTTTAGTATACATGCATTTACTTATAATACAGTTCGGTGTCGGTCTCGTA
GNNNNNNNNNNNNNNNNNNNNATCGATGAATTCGAGCTCGTTTTCGACACTGGATG
GCGGCGTTNNNNNttagttttgctggccgcatc

For *OMN773* and *OMN775* a stretch of 5 random nucleotides (N5) was added after the stop codon to include a clone barcode. Additionally, to generate new barcoded WT strains, OMN775 contains an additional stretch of 20 nt.

The PCR product resulting from *OMN761-OMN773* contains homology to regions within the original deletion cassette targeting the junction of the TEF1-KAN and the 1 nt upstream of the genotype barcode (D2-Downtag-D1), therefore shortening the terminator (from 262 nt in the original YKOC to 43 in the RNA-barcoded collection) enabling the use of the endogenous terminator. A total of 7 different WT strains were generated with distinct clone and genotype barcodes as controls. These wild type strains were verified by Sanger sequencing. For wild type strains the PCR product resulting from *OMN774-775* contains homology regions upstream and downstream of the URA3 loci.

Generation of the PCR cassette was done by pairing *OMN761-OMN773* and *OMN774-OMN775* for deletion strains and WT strains respectively. Expand High Fidelity PCR System (Roche, 11759078001) was used to amplify the *URA3* marker using 10 ng pRS406 as a template in reactions of 100 µl (Buffer# 2 10X with $MgCl_2$, dNTPs mix 1 mM (Promega, U1420), 1µM OMN761 or OMN773 (Fw), 1 µM of OMN773 or OMN774, and 2.6 U (0.75 µl) of Expand High Fidelity Enzyme Mix. PCR product was purified using columns (Qiagen, 19066), transferred to a DNA purification column (EconoSpin Columns, 1910-050) and centrifuged for 1 minute at 13000 rpm, washed once with PE Buffer (Quiagen, 19065) and eluted using nuclease free $H_2O$ (centrifuge 1 min 3000 RPM). Finally, the purified PCR product was diluted to 400 ng/µl.

**Sequence of the RNA-barcoded Yeast Knock Out deletion structure.**

GATGTCCACGAGGTCTCTNNNNNNNNNNNNNNNNCGTACGCTGCAGGTCGACGGATCCCCGGGT
TAATTAAGGCGCGCCAGATCTGTTTAGCTTGCCTCGTCCCCGCCGGGTCACCCGGCCAGCGACA
TGGAGGCCCAGAATACCCTCCTTGACAGTCTTGACGTGCGCAGCTCAGGGGCATGATGTGACTG
TCGCCCGTACATTTAGCCCATACATCCCCATGTATAATCATTTGCATCCATACATTTTGATGGC
CGCACGGCGCGAAGCAAAAATTACGGCTCCTCGCTGCAGACCTGCGAGCAGGGAAACGCTCCCC
TCACAGACGCGTTGAATTGTCCCCACGCCGCGCCCCTGTAGAGAAATATAAAAGGTTAGGATTT
GCCACTGAGGTTCTTCTTTCATATACTTCCTTTTAAAATCTTGCTAGGATACAGTTCTCACATC
ACATCCGAACATAAACAACC**atgggtaaggaa**tcgaaagctacatataaggaacgtgctgctac
tcatcctagtcctgttgctgccaagctatttaatatcatgcacgaaaagcaaacaaacttgtgt
gcttcattggatgttcgtaccaccaaggaattactggagttagttgaagcattaggtcccaaaa
tttgtttactaaaaacacatgtggatatcttgactgattttccatggagggcacagttaagcc
gctaaaggcattatccgccaagtacaatttttactcttcgaagacagaaatttgctgacatt
ggtaatacagtcaaattgcagtactctgcgggtgtatacagaatagcagaatgggcagacatta
cgaatgcacacggtgtggtgggcccaggtattgttagcggtttgaagcaggcggcggaagaagt
aacaaaggaacctagaggccttttgatgttagcagaattgtcatgcaagggctccctagctact
ggagaatatactaagggtactgttgacattgcgaagagcgacaaagattttgttatcggcttta
ttgctcaaagagacatgggtggaagagatgaaggttacgattggttgattatgacacccggtgt
gggtttagatgacaagggagacgcattgggtcaacagtatagaaccgtggatgatgtggtctct
acaggatctgacattattattgttggaagaggactatttgcaaagggaagggatgctaaggtag
agggtgaacGTTACAGaaaagcaggctgggaagcatatttgagaagatgcggccagcaaaacta
aNNNNNAACGCCGCCATCCAGTGTCGAAAACGAGCTCGAATTCATCGATNNNNNNNNNNNNNNN
NNNNNCTACGAGACCGACACCG

Sequence legend:

N: Random nucleotides N20 genotype barcodes from original YKOC and N5 represents the newly added clone barcode.
Lowercase: open reading frame of the URA3 gene. Bold letters denote nucleotides remaining from the original G418 resistance. These nucleotides were kept to increase the efficiency of the integration.
Underlined regions: original D2 and D1 sequences from the original YKOC

**Modification of the Yeast Knock Out Collection.**
For the generation of the modified strains, frozen glycerol stocks from the haploid yeast knock out collection were grown on YPD (Yeast Peptone Dextrose medium) supplemented with G418 (Geneticin, 200 mg/L) into 96 well plates using a Robot (Singer instruments). Strains were allowed to grow for 48 hours until saturation at 25ºC before transformation.

For high-throughput liquid transformation, 10 µl of saturated cultures were transferred and diluted into deep 96 well plates containing 700 µl of YPD using EpMotion 96 (Eppendorf). Cells were allowed to grow for 6 hours, media removed and 200 µl of LiAc solution was dispensed using a Multidrop Combi (Thermo, 5840300). A 1:1 mix of 40 µl of ssDNA (10 mg/ml) and a denatured PCR cassette for deletion strains (PCR product OMN761-773) and for WT strain (OMN774-OMN775) (400 ng/µl) was added to the cell-LiAC mix with an EpMotion. Last, 300 µl of 50% PEG solution was added to the mix. Yeast transformation plates were incubated at 30ºC for 1 hour and after 35 µl of DMSO was dispensed and briefly shacked with Multidrop Combi. Transformations were heat shocked for 30 minutes at 42ºC using a water bath, cells were centrifuged, and transformation reagents removed. Pelleted cells were washed with 200 drop out media (MPBio, 1145112-CF) centrifuged, resuspended in 300 µl of URA- and incubated at 30ºC overnight. Then plates were centrifuged, and cells resuspended in fresh URA- media for 48 hours. Transformation efficiency was determined by transferring 10 µl of the transformation into fresh URA- media and check for growth using a microplate reader (Synergy HX1, Agilent Technologies). Grown cells were mixed with URA- 50% glycerol to make frozen stocks of the new transcriptionally barcoded Yeast Knockout Collection. The genotype genomic location of reengineered strains was validated using the 3'UTR region of URA3 and the median distance to the expected deletion has can be found in Extended Data Table 1.

**Yeast Growth and harvest for the Perturb-seq experiments.**
Frozen glycerol stocks of each individual mutant from the transcriptionally barcoded yeast knock out collection were recovered on URA- media at 25ºC for 48 hours until saturation. The next day, 5 µl of cultures were refreshed into a new 96 well plate containing 200 µl of YPD. Cells were allowed to grow for 6 hours until they reached mid exponential phase average OD660 0.6-0.8. To maximize representativeness of each mutations optical density for all plates was assessed after seeding and before pooling using a Synergy HX1 reader. Cells then were pooled together into large flasks and shacked vigorously to ensure a homogeneous mixture.
We subjected or not cell pools to osmostress (0.4M NaCl for 15 minutes) and followed the cell fixation protocol from GEXSCOPE® Microbial Single Cell RNA Library Kit HD (Yeast) (4161031). The selection of this experimental conditions is based on a combination of extensive transcriptomics data [2,3] that determines the peak of expression of osmoresponsive genes. Briefly, 100 ml cells where centrifuged 1 min 3000 RPM, media poured off and cells immediately fixed and resuspended in 10 ml of ice cold 80% Methanol (Scharlab, ME0301005P). For each condition methanol-fixed cells were split into 20 different aliquots (1 ml) from each condition (control/NaCl) and stored at -20ºC, to avoid multiple freezing and thawing.

**Library generation.**
To perform the yeast, Perturb-seq experiments we followed GEXSCOPE® Microbial Single Cell RNA Library Kit HD instructions (4161042, Singleron Biotechnologies). Briefly, A methanol frozen aliquot of each condition was equilibrated for 15 minutes at 4ºC (ice) before pelleted (2 min 1640 g at 4ºC). Supernatant was discarded and cells were washed twice with 700 µl of rehydration buffer (DPBS1X, BSA 20 mg/ml (Thermo Fisher Scientific, 14190144 and AM2616), RNASe inhibitor (40U/µl) (Takara, 2313A), Actinomycin D (2mg/ml) (Sigma, A1410-2MG)). To disaggregate potential cell clumps we filtered twice with PluriStainer (PluriSelect,43-50040-03) and cells were counted with a Neubauer chamber. A total of 220,000 cells (120 µl) of the yeast suspension per matrix-cartridge in the High Density Singleron Matrix v1.0.1instrument. For each

run, a total of 2 matrix-cartridges were run simultaneously each one loaded with control and NaCl simultaneously. The position of each sample in the Singleron Matrix instrument (upper/lower slot) was exchanged per every run. A total of 29 matrix-cartridges were run (14 control and 15 NaCl). Single-Yeast Partitioning, mRNA capture, reverse transcription, cDNA amplification and cDNA purification were performed according to manufacturer's instructions. For cDNA library amplification 8 PCR cycles were used. Samples were processed per run (as group of two samples, control/NaCl) until cDNA purification. The cDNA quality of each cDNA was assessed by Qubit (Q33231, Thermo Fisher Scientific, Qubit™ 1X dsDNA HS Assay Kits), the size and integrity of the full-length cDNA was inspected with a DNA Pico Bioanalyzer chip (Agilent technologies). Library preparation was done using 50 ng of purified cDNA as an input and library amplification was done with 10 PCR cycles. Finally, library quality check was performed with Qubit and library size was determined with DNA Pico Bioanalyzer chip. Equimolar pools of the 29 libraries were pooled together and sequenced in one S4 NovaSeq lane (CeGAT, Tübingen) using paired end 150 cycles (PEx150 cycles). A total of 3.58 Tb of data were generated from a single run.

**Targeted Amplification (TA-libraries).**

For each of the 29 full length cDNA libraries, a total of 5ng was used as an input to amplify the *URA3* transcript and the 3'UTR with a one-step PCR reaction. PCR was generated using the NEBNext Ultra II (E7645L, New England Biolabs) in a final volume of 25 µl following manufacturer's instructions. The primer design incorporates the P5/P7 sequences, the index barcode and the Illumina read 1/2 sequences to enable direct sequencing of the purified PCR products. The resulting PCR products (102 bp excluding the primer overhangs) were purified using Ampure Beads at a 1.5X ratio and eluted with 30 µl of elution buffer. Library size was inspected by Bioanalyzer chip and equimolar amounts of each library were pooled together for sequencing using a NextSeq500 (Illumina).

**Data analyses.**

All the code generated from the scRNA-seq data has been uploaded to GitHub. This code contains the analysis pipeline and the code to reproduce the figures. The fully processed R object containing both Perturb-seqs combined or separated into individual objects with the corresponding metadata has been uploaded with the raw sequencing data.

**Read pre-processing, alignment, and filtering**.

To process the sequencing data (FASTQ), we used standard CeleScope (v1.14) pipeline from Singleron (https://github.com/singleron-RD/CeleScope). To process FATSQ files, we first trimmed the D1+downstream and D2+upstream reads originating from the knock-in loci to retain only the genotype sequences. Reads were aligned to a Yeast's sacCer3 reference genome with all the genotype sequences appended as additional contigs to obtain counts of gene and genotype barcode expressions. Sequences of the genotype barcode were downloaded from the Yeast Deletion Project (http://www-deletion.stanford.edu/YDPM/YDPM_index.html). Additionally, an artificial chromosome containing the clone barcodes (5 nucleotides), the common terminator region and the genotype barcode (D2-Downtag-D1) was added to the reference genome to enable genotype identification from the expression matrix. Example of an artificial genotype chromosome:

>bc-Systematic Name
NNNNNAACGCCGCCATCCAGTGTCGAAAACGAGCTCGAATTCATCGATNNNNNNNNN
NNNNNNNNNNNNNCTACGAGACCGACACCG

Naming of replacement strains; the original YKOC and commercial collections contain some replacement strains which represent repetitions for conflictive strains. The names of these replacement strains and genotype barcodes are identical between repetitions. To include these strains in the reference genome and to avoid a naming conflict, chromosomes of each repeated strain was named sequentially: bc-systematic name, bc-systematic name-2, or bc-systematic name-3.

**Analysis of the targeted amplicon library**

Barcode and UMI information were extracted from FastQ files with Singleron Celescope software version 1.14.1, using commands 'celescope rna sample --chemistry auto' and 'celescope rna barcode', and Singleron v3 whitelist and linker files.

Resulting fastq files were imported into R. Clone information was extracted from the sequenced read at positions 21 to 25. In order to find the genotype information, the region between the D1 and D2 inserts was extracted by matching the corresponding sequences allowing at most 3 mismatches. Only reads containing both D1 and D2 were kept for further analysis. In case of multiple matches of any of the insert sequences the one with maximum start position was chosen. Reads were further filtered for starting matching positions of D1 between 85 and 90, and D2 between 48 and 51. The inserted sequence corresponding to knock out genotypes in the library were compared to the remaining sequences using the vmatchPattern R function with a maximum mismatch of 2. Cells assigned to more than one genotype with no genotype present in more than 70% of reads were discarded.

To combine both sets of assignments, cells with different genotypes assigned in both libraries were marked as "conflicted" and omitted from downstream analysis. Cells with the same genotype assigned in both libraries were given that genotype. Cells with genotype assigned in only one of the 2 libraries were also given the assigned genotype.

**Genotype position.**

Because the YKOC is a globally used resource the assessment of the location of each deletion has only been performed manually for selected mutants or only available for the homozygote diploid collection at a global scale with Whole Genome Sequencing [4] . Our strategy enables to define the genomic location of each deletion in the new KO collection by leveraging the high resolution of the targeted amplification which has coverage of the endogenous terminator. To assign the genomic position all the FASTQ reads were combined and aligned against the yeast genome (saccer3, SGD). The median nucleotide distance between the mapped read and the annotated STOP codon was done to assess the genomic position of the intended deletion. Genotypes with greater distances greater than 300 bp from the genomic loci were considered incorrect (Extended Data Table 1).

**Downstream Processing.**

The complete sc-RNA Yeast Genome Dataset includes a single processed Seurat object (Seurat v4,) [5] that contains all cells profiled in both conditions and the corresponding metadata. Additionally, we generated an additional Seurat object corresponding to the control or stressed samples and the corresponding metadata available at E-MTAB-14004. As it has been done before and to ease the information to the community given the wide usage of the YKOC we have reported all the information for all genotypes detected and removed incorrect genotypes for detailed analysis and experimental validation.

To generate the Seurat Objects, the outputs of Celescope were used to generate the corresponding cell expression matrix either by combining both conditions or for each condition individually. To normalize gene expression across cells, we applied the SCTransform procedure and regressed out the cell cycle scores by supplying the cell cycle variable genes to the vars.to.regress argument. The final log normalized results were used for all downstream analysis. We followed standard Seurat clustering guidelines with the following parameters and modifications: We calculated cell cycle scores for each cell by scoring the cell cycle signature using canonical cell cycle genes previously used in scRNA-seq [6] (see gene lists, Extended Data Table 7) by using the CellCycleScoring function. We also identified highly variable genes using the FindVariableFeatures from the Seurat package with nfeatures =1. To perform cell clustering, first we performed a linear dimensional reduction using the "RunPCA" function from Seurat Package using PC1 and PC2. Visualization of gene loadings for the complete dataset was done by using the VizDimLoading function of PC1. To cluster cells we then applied the Seurat pipeline FindNeighbors (dims 1:14) and FindClusters (resolution =1). To visualize the UMAPs we used RunUMAP (dims 1:14).

**Clone comparison.**

Only the 220 genotypes with more than 200 cells were considered for the analysis. For each genotype, clones with more than 9 cells were compared against the rest of the clones (with at least 3 cells) using the FindMarkers function from Seurat (adjusted *pvalue* < 0.05).

**Cell state markers.**

To extract cell state markers we applied the differential expression function included in Seurat through FindAllMarkers for the complete dataset (both conditions) and each condition individually. Gene ontology enrichments of upregulated cell state markers were performed using Metascape v3.5.20230501 [7]default parameters and we used S. *cerevisiae* as input and output specie. The enrichment terms per each input list was downloaded and appended to Extended Data 2 and 3.

To visualize the expression signatures per each condition individually, we generated lists of the all genes upregulated genes of each cluster using the FeaturePlot (order=TRUE). To visualize the co-expression of the aging signature we used the UCell package AddModuleScore_UCell aging genes defined by RNA-seq. The top 15 aging genes were retrieved from Patnaik et al (https://pubmed.ncbi.nlm.nih.gov/35858543/) [8], and used to generate an unbiased cell signature, or either a signature split in iron-containing or non-iron signature. The gene list of induced Environmental Response genes was obtained from published datasets [9,10] using the updated list from Gasch et al., 2017 (https://sgd-prod-upload.s3.amazonaws.com/S000343511/ESR_clusters_UPDATED_2017.xlsx). Gene lists used for signatures are listed in Extended Data Table 7.

**Conservation cell state enrichment across conditions.**

To calculate the degree of similarities between states, we calculated the correlation of expression between the upregulated cell state markers across clusters identified in each condition independently. To calculate the degree of similarities between states, a Fisher Test was performed comparing the common upregulated markers (adjusted *pvalue* < 0.05 and average log2 Fold Change > 0.25) between conditions cell states.

**Cell state genotype enrichment.**

Cell state enrichment per genotype was done for genotypes with ≥6 cells to avoid biases due to cell number. To determine the enrichment degree a Fisher Test was performed and the odds ratio of each genotype per cluster was calculated. We considered enriched genotypes if the odds ratio was >1 and adjusted *pvalue* <0.05 or as depleted if the odds ratio was <0 and adjusted *pvalue* < 0.05. Cell state enrichment per each genotype is reported in Extended Data Table 4 and 5 respectively.

To visualize the protein interaction between enriched genotypes in a specific cluster we used the Metascape v3.5.20230501 default parameters. These parameters only include physical interactions using STRING (physical score > 0.132) and BioGrid. Additionally Molecular Complex Detection (MCODE) algorithm was used to define densely connected networks within the Metascape default parameter app.

**Leverage Score (transcriptional heterogeneity).**

Differential gene expression was assessed using the Wilcoxon rank sum test. Each mutant was compared to wild type in control and stress samples separately. For each comparison, genes with 0 counts were first removed. To optimize statistical power, independent filtering was applied based on mean normalized expression. To identify the optimal threshold for average counts, we: 1) Applied different thresholds to filter out genes with averaged counts less than the thresholds. 2) Performed *pvalue* adjustment using Benjamini-Hochberg procedure on the remaining genes that passed the threshold and 3) Counted the number of statistically significant genes with adjusted p values lower than 0.05. The threshold resulting in the highest number of significant genes was finally used. This threshold was found and applied separately for the control (<1.27) and stress treated samples (<1.19).

The leverage score was calculated in the same way as in [11], except that we used log normalized counts yielded from SCtransform procedure instead of plate level z-score because of the variable composition of the wild type cells in each plate that might introduce more variance. Specifically, we 1) Constructed a count matrix depicting cells in rows and genes in columns, consisting of all genes with mean expression >0.25 UMI counts per cell. 2) We calculated the top 20 left singular vectors for each plate using the partial SVD algorithm (scipy.sparse.linalg.svds) with the arpack solver and k=20. Row wise square norm of the resulting *n* by 20 matrices (where *n* is the number of cells) was calculated to give the leverage scores. These scores were normalized so that the sum over all cells in each plate is 1. 3) To normalize leverage scores across plates, we log transformed the scores from the previous step, and z-score normalized these scores relative to the scores of wild type cells (subtracting the mean and dividing by the standard deviation of the wild type cells). We reported the leverage score and the scaled leverage score of each cell in the corresponding metadata. Extended Data 7 contain the average, standard deviation, variance of the leverage score raw and scaled per each condition.

We ranked genotypes according to the standard deviation of the leverage score (wild type =1). For both conditions, and similar to the cell state enrichment in Fig. 3, and only for genotypes with at least 6 cells (threshold defined in Extended Data Fig. 4a, 4b). The standard deviation was calculated using base R functions and the dplyr package[12]. We then visualized the negative drivers (≥1.3 standard deviation leverage score) or positive drivers (≤ 0.7 standard deviation leverage score) per each condition. The projection into the yeast interaction network was done using TheCellMap using the overlay function with the default parameters (https://thecellmap.org/) [13]. Functional enrichment analysis of control or stress genotypes with increased (>30%) or decreased heterogeneity (<30% of the wild type strain) was analyzed the Spatial Analysis of Functional Enrichment (SAFE) build in TheCellMap (www.thecellmap.org), applying default settings.

### Experimental validations

#### Cell state reporters.

Recombinant DNA techniques and transformation of bacterial and yeast cells were performed using standard methods. To generate reporters for cell states, we used the MoClo Yeast Toolkit Modular cloning system [14]. Building of the plasmid constructs was achieved using Golden Gate assembly. Each reporter contains a transcription unit composed of: the corresponding promoter (700 bp upstream of the annotated ATG), UbiM degradation signal, florescent protein and terminator (300 bp downstream of annotated STOP codon). All new part sequences were either mutated or synthesized to avoid of the BsmBI, BsaI, and NotI recognition sequences. Promoter and terminator sequences were amplified from BY4741 genomic DNA and purified using the MiniElute PCR purification (28004, Quiagen). Plasmids generated in this study are described in Table 1.
Entry plasmids were generated using the MoClo guidelines and were amplified in *Escherichia coli* DH5α competent cells grown at 37ºC in LB medium supplemented with the corresponding antibiotic for selection. Plasmid extraction was done using the E.Z.N.A.® I Kit (D6942-02, Avantor) and verified by Sanger sequencing. Purified plasmid was linearized with NotI (NEB) and integrated into the yeast genome. Standard yeast transformation was done using the LiAc method into the corresponding yeast background and colonies were selected by marker selection and colony PCR. Strains harboring cell state reporters generated in this study are described in Table 2. Expression of the reporters was followed by flow cytometry (see below).

#### Calcofluor white staining and mitochondrial morphology (bud scars; aging phenotype analyses).

Wild type cell or cells carrying the corresponding reporter, were grown to exponential phase in SC media. Cells were filtered with a 70 µm mesh and a total of 300,000 cells was sorted using a FACS Aria III cell sorter into 15 ml falcons. The top 2% of the population (*pFIT3*-mCherry and *pHUG1*-mCherry) were fixed by directly sorting into Ethanol 100%. Additionally, a random sort for the entire population or for a wild type strain were collected as controls using the same fixation strategy. Fixed cells were stored at 4ºC until Calcofluor white staining.
To visualize and count bud scars, we stained cells with 200 µg/ml Calcofluor White Stain as reported [15] (18909 Fluka Analytica) with minor modifications. Briefly, the indicated populations of cells were sorted directly into Ethanol fixed cells and washed once with DPBS 1X (J67653.K2, Thermo Fisher) to remove excess Calcofluor. Scars per each cell for each corresponding population were counted using at 100x magnification (Plan Apo VC 60x Oil objective) using a Nikon Eclipse Ti inverted microscope and an ORCA digital camera (Hamamatsu). Per each biological replicate a total of 250-300 cells were counted.

To assess mitochondrial morphology, we used the same procedure as above except that cells were sorted into rich media (YPD) containing 100 nM MitoTracker™ Red CMXRos (Invitrogen, M7512). Cells were incubated in this media for 30 minutes, washed with media without MitoTracker and fixed in YPD 4% formaldehyde and stored for imaging. The morphology of mitochondria was assessed for at least cells per each biological replicate of the indicated strains.

**Chromatin Immunoprecipitation.**
Cells were grown to mid exponential log phase and 50 ml were harvested at OD660 0.6. Fixation was done by adding 1% formaldehyde (Sigma, F1635) for 20 minutes and quenched with Glycine 125 mM for 15 minutes at room temperature. Cells were pelleted by centrifugation and washed with TBS 1X four times at 4ºC. Chromatin immunoprecipitation was done as described in [16] Briefly, a total of 0.5 ug antibody per sample against Lysine acetylation (Cell Signaling, 9441S) and conjugated to 25 µ rabbit Dynabeads® M-280 Sheep Anti-Rabbit IgG (Life Technologies, 11204D) overnight at 4ºC. Levels of Lysine acetylation was determined by qPCR using primers specific to the indicated promoters to: PHO84 (*Fw: GGACGTGTTATTTCCAGCAC and Rv: CAGGCAAAACGGGAGAAGAG*) and VTC1 (*Fw: TTGGCATCGCTATTTTCGGA and Rv: ACCGACCGTAACAAGCGATA*) and as loading control an intergenic region of the right arm of choromosome VI was used (*Fw: ACCACTCAAAGAGAAATTTACTGGAAGA and Rv: CTCGTTAGGATCACGTTCGAATC*). For qPCR Power SYBR™ Green PCR Master Mix (Life Technologies, 4368708) was used in a final volume of 10 µl following manufacturer's protocol and qPCR reaction conditions. The resulting Ct valies each gene was normalized to the loading control and delta Ct method [17] as used to calculated abundance of acetylated Lysines. The values for the wild type strain set to 1 and used as a reference.

**Competition assays.**
Wild type strains carrying the indicated expression reporters were grown to mid exponential log phase and sorted using Aria SORP (Becton Dikinson). A total of 20,000 cells of each top 2% of the population was sorted and 20,000 cells of a wild type strain carrying a constitutive GFP (*pTEF1*) was sorted on top in a final volume of 200 µl of rich media (YPD). A total of 150 µl of the mixed culture was fixed with 4% formaldehyde as time 0. The remaining culture was evenly split into YPD and YPD 1M NaCl. Cells were diluted every 24 hours and fixed after 48 hours from t0.

**Flow cytometry analysis.**
Cells were recorded from each sample according to their FSC and SSC distributions and unmixed to identify the fluorescence signal for each fluorophore (mCherry, iRFP or GFP). For competition assays, cells were gated based on the constitutive expression of GFP of the wild type strain versus the side scatter for three biological replicates. To read the expression of pFIT3-mCherry reporter in mutants in mutants enriched in cluster 8C, the full spectrum of 10,000 cells were recorded Cytek® Aurora (4-laser and 64 Fluorescence Emission Detection Channels) gated according to the FCS and SSC distributions. The unmixed signal was used to assess the expression distribution of each mutant against the wild type and the mean expression (arbitrary units) per each strain and biological triplicates. The mean and median expression of each strain was obtained for 3 biological replicates. Cytometry data were analyzed using FlowJo™ Software (BD Life Sciences).

**Comparison of fitness scores.**

This analysis was performed using Supplementary Table 1 (Mutant Fitness Conditions) from https://pubmed.ncbi.nlm.nih.gov/33958448/, encompassing 14 stressing conditions and 4429 genotypes. Only common genotypes between the publicly available dataset and our data were included. Three gene sets were built: the first comprised the top 50 genotypes with the highest standard deviation of the scaled leverage score, the second consisted of the bottom 50 genotypes with the lowest standard deviation of the scaled leverage score, and the third, called biased, that comprised enriched genotypes (adjusted *pvalue* < 0.05 and an odds ratio > 1) identified through the Fisher test from Section. A permutation test was conducted for each stress score and each geneset by comparing the mean of the gene set with the mean of 1000 randomly selected unclassified genotypes of equal length.

**Growth curves.**

The indicated strains were grown below $OD_{(660)}$ =1 in rich media. For the experiment cells were washed three times with complete synthetic media pH=5.5 or pH=8.8. Then cells were diluted to $OD_{(660)}$=0.05 in a final volume of 200 µl in a 96 well plate. Plates were incubated under orbital shaking at 30°C in a Synergy H1 (BioTek® Instruments) and $OD_{(660)}$ was recorded every 30 minutes for 48 hours. The Lag Time was calculated using the Gene 5 software((BioTek® Instruments).

**Table 1. List of plasmids used for experimental valudation**

| Plasmid num | Plasmid Name | Description | Source |
|---|---|---|---|
| pDC198 | pYTK-Spect-*LEU2*_int-*HIS3* (pYTK147) | backbone for integrative plasmids | [18] |
| pNO82 | pYTK-Nat-HO_int-Nat (pYTK168) | backbone for integrative plasmids | This study |
| pRP093 | pYTK147-*pFIT3*-UbiM-mCherry 3b-*tFIT3* | Integrating plasmid carrying the *FIT3* reporter consisting of the promoter of *FIT3*, a n N-terminal degradation signal fused to mCherry and terminator *FIT3*. | This study |
| pRP108 | pYTK147-pHUG1 dBsaI-UbiM-mCherry 3b-tHUG1 | Integrating plasmid carrying the *HUG1* reporter consisting of the promoter of HUG1, a n N-terminal degradation signal fused to mCherry and terminator *HUG1*. dBsaI indicates the internal BsaI site was mutated to enable correct assembly. | This study |
| pRP128 | pYTK168-pHSP12-UbiM-IRFP 3b-tHSP12 | Integrating plasmid carrying the *HSP12* reporter consisting of the promoter of HSP12, a n N-terminal degradation signal fused to mCherry and terminator *HSP12*. | This study |

**Table 2. List of strains generated in this study for experimental validation**

| Strain name | Genotype | Source |
|---|---|---|
| YMN75 | *ENO1*-mCherry::HPH | This study |
| yRP067 | *ENO1*-iRFP::NAT (YCS368) (Clone 3) | This study |
| yRP145 | BY4741-pYTK147-pFIT3-UbiM-IRFP 3b- tFIT3:HIS | This study |
| yRP148 | BY4741-pYTK147-pFIT3-UbiM-mCherry 3b- tFIT3:HIS | This study |
| yRP171 | BY4741-pYTK147-pHUG1 dBsaI-UbiM-mCherry 3b- tHUG1:HIS | This study |
| yRP175 | BY4741-pYTK147-pTMA10 -UbiM-mCherry 3b- tTMA10:HIS | This study |
| yRP178 | BY4741-pYTK144-TEF1i-YmukG1-tTDH1:LEU | This study |
| yRP193 | BY4741-pYTK168-pHSP12-UbiM-IRFP 3b-tHSP12: NAT | This study |
| yRP195 | BY4741-pYTK147-pSBH2 dBsaI-UbiM-mCherry 3b- tSBH2:HIS | This study |
| yRP201 | EAF3::URA3-pYTK147-pFIT3-UbiM-mCherry-tFIT3:HIS | This study |
| yRP204 | GRX5::URA3-pYTK147-pFIT3-UbiM-mCherry-tFIT3:HIS | This study |
| yRP206 | KCS1::URA3-pYTK147-pFIT3-UbiM-mCherry-tFIT3:HIS | This study |
| yRP208 | MRH4::URA3-pYTK147-pFIT3-UbiM-mCherry-tFIT3:HIS | This study |
| yRP211 | MRP7::URA3-pYTK147-pFIT3-UbiM-mCherry-tFIT3:HIS | This study |
| yRP213 | MRPL49::URA3 (3_I)-pYTK147-pFIT3-UbiM-mCherry-tFIT3:HIS | This study |
| yRP216 | MRPL49::URA3 (4_N)-pYTK147-pFIT3-UbiM-mCherry-tFIT3:HIS | This study |
| yRP219 | MRPL51::URA3-pYTK147-pFIT3-UbiM-mCherry-tFIT3:HIS | This study |
| yRP220 | MRPL7::URA3-pYTK147-pFIT3-UbiM-mCherry-tFIT3:HIS | This study |
| yRP223 | MRPS16::URA3-pYTK147-pFIT3-UbiM-mCherry-tFIT3:HIS | This study |
| yRP225 | MSS116::URA3-pYTK147-pFIT3-UbiM-mCherry-tFIT3:HIS | This study |
| yRP227 | PHO85::URA3-pYTK147-pFIT3-UbiM-mCherry-tFIT3:HIS | This study |
| yRP230 | RIM101::URA3-pYTK147-pFIT3-UbiM-mCherry-tFIT3:HIS | This study |
| yRP231 | RIM8::URA3 (4_G)-pYTK147-pFIT3-UbiM-mCherry-tFIT3:HIS | This study |
| yRP234 | RIM8::URA3 (4_N)-pYTK147-pFIT3-UbiM-mCherry-tFIT3:HIS | This study |
| yRP238 | SAS4::URA3-pYTK147-pFIT3-UbiM-mCherry-tFIT3:HIS | This study |
| yRP240 | SAS5::URA3-pYTK147-pFIT3-UbiM-mCherry-tFIT3:HIS | This study |
| yRP241 | SNF7::URA3-pYTK147-pFIT3-UbiM-mCherry-tFIT3:HIS | This study |
| yRP244 | SNF8::URA3-pYTK147-pFIT3-UbiM-mCherry-tFIT3:HIS | This study |
| yRP245 | SNT1::URA3-pYTK147-pFIT3-UbiM-mCherry-tFIT3:HIS | This study |
| yRP248 | SOV1::URA3-pYTK147-pFIT3-UbiM-mCherry-tFIT3:HIS | This study |
| yRP251 | SUR1::URA3-pYTK147-pFIT3-UbiM-mCherry-tFIT3:HIS | This study |
| yRP253 | VPS28::URA3-pYTK147-pFIT3-UbiM-mCherry-tFIT3:HIS | This study |
| yRP256 | VPS36::URA3-pYTK147-pFIT3-UbiM-mCherry-tFIT3:HIS | This study |
| yRP258 | YGR122W::URA3-pYTK147-pFIT3-UbiM-mCherry-tFIT3:HIS | This study |
| yRP261 | YMN478-pYTK147-pFIT3-UbiM-mCherry-tFIT3 | This study |
| yRP264 | YMN479-pYTK147-pFIT3-UbiM-mCherry-tFIT3 | This study |
| yRP266 | YPL216W::URA3-pYTK147-pFIT3-UbiM-mCherry-tFIT3:HIS | This study |

**Methods References**


1. Winzeler, E. A. *et al.* Functional characterization of the S. cerevisiae genome by gene deletion and parallel analysis. *Science (1979)* **285**, 901–906 (1999).
2. Nadal-Ribelles, M. *et al.* Hog1 bypasses stress-mediated down-regulation of transcription by RNA polymerase II redistribution and chromatin remodeling. *Genome Biol* **13**, R106 (2012).
3. Latorre, P. *et al.* Data-driven identification of inherent features of eukaryotic stress-responsive genes. *NAR Genom Bioinform* **4**, (2022).
4. Puddu, F. *et al.* Genome architecture and stability in the Saccharomyces cerevisiae knockout collection. *Nature* vol. 573 416–420 Preprint at https://doi.org/10.1038/s41586-019-1549-9 (2019).
5. Hao, Y. *et al.* Integrated analysis of multimodal single-cell data. *Cell* **184**, 3573-3587.e29 (2021).
6. Jackson, C. A., Castro, D. M., Saldi, G. A., Bonneau, R. & Gresham, D. Gene regulatory network reconstruction using single-cell rna sequencing of barcoded genotypes in diverse environments. *Elife* **9**, (2020).
7. Zhou, Y. *et al.* Metascape provides a biologist-oriented resource for the analysis of systems-level datasets. *Nature Communications 2019 10:1* **10**, 1–10 (2019).
8. Andreatta, M. & Carmona, S. J. UCell: robust and scalable single-cell gene signature scoring. doi:10.1101/2021.04.13.439670.
9. Gasch, a P. *et al.* Genomic expression programs in the response of yeast cells to environmental changes. *Mol Biol Cell* **11**, 4241–57 (2000).
10. Gasch, A. P. *et al.* Single-cell RNA sequencing reveals intrinsic and extrinsic regulatory heterogeneity in yeast responding to stress. *PLoS Biol* **15**, (2017).
11. Replogle, J. M. *et al.* Mapping information-rich genotype-phenotype landscapes with genome-scale Perturb-seq. *Cell* **185**, 2559-2575.e28 (2022).
12. GitHub - tidyverse/dplyr: dplyr: A grammar of data manipulation. https://github.com/tidyverse/dplyr.
13. Usaj, M. *et al.* TheCellMap.org: A web-accessible database for visualizing and mining the global yeast genetic interaction network. *G3: Genes, Genomes, Genetics* **7**, 1539–1549 (2017).
14. Lee, M. E., DeLoache, W. C., Cervantes, B. & Dueber, J. E. A Highly Characterized Yeast Toolkit for Modular, Multipart Assembly. *ACS Synth Biol* **4**, 975–986 (2015).
15. Zahner, J. E., Harkins, H. A. & Pringle, J. R. Genetic analysis of the bipolar pattern of bud site selection in the yeast Saccharomyces cerevisiae. *Mol Cell Biol* **16**, 1857 (1996).
16. Solé, C. *et al.* Control of Ubp3 ubiquitin protease activity by the Hog1 SAPK modulates transcription upon osmostress. *EMBO J* **30**, 3274–3284 (2011).
17. Livak, K. J. & Schmittgen, T. D. Analysis of relative gene expression data using real-time quantitative PCR and the 2(-Delta Delta C(T)) Method. *Methods* **25**, 402–408 (2001).
18. Canadell, D. *et al.* Implementing re-configurable biological computation with distributed multicellular consortia. *Nucleic Acids Res* **1**, 1–18 (2022).
